# Human bone marrow mesenchymal stem/stromal cell behaviour is coordinated via mechanically activated osteocyte-derived extracellular vesicles

**DOI:** 10.1101/730077

**Authors:** Kian F. Eichholz, Ian Woods, Gillian P. Johnson, Nian Shen, Michele Corrigan, Marie-Noelle Labour, Kieran Wynne, Michelle C. Lowry, Lorraine O’Driscoll, David A. Hoey

## Abstract

Osteocytes are mechanosensitive cells that are believed to play a fundamental role in coordinating bone mechanoadaptation via the secretion of paracrine factors. However, the exact mechanisms by which osteocytes relay mechanical signals to effector cells is poorly understood. In this study, we demonstrated that osteocytes subjected to a physiologic fluid shear secrete a distinct collection of factors that significantly enhance human MSC recruitment and osteogenesis. Utilising proteomics we generated an extensive map of proteins within the mechanically activated osteocyte secretome, identifying numerous paracrine factors that are modified by mechanical stimulation. Moreover, we identified the presence of extracellular vesicles (EVs) and further demonstrated that these mechanically activated osteocyte derived EVs (MAEVs) coordinate human MSCs recruitment and osteogenesis. This indicates that mechanical conditioning of parent cells can modify EVs and demonstrates the pro-osteogenic potential of MAEVs as a cell-free therapy to enhance bone regeneration and repair in diseases such as osteoporosis.

## Introduction

Osteocytes are the most abundant cell type in bone and are known as the primary sensing and metabolism-controlling cells within the tissue. Osteocytes are key to directing the processes of bone formation and resorption via the secretion of various signalling factors which act upon bone forming osteoblasts and resorbing osteoclasts and their progenitors, skeletal and haematopoietic stem cells [1]. The implications of this can be seen in the highly debilitating and life threatening disease that is osteoporosis, which has been linked to osteocyte apoptosis [2] and reduced osteocyte numbers in affected patients [3]. This results in a significant drop in quality of life, increased risk of additional complications due to immobilisation, and significantly increased mortality rates due to fracture and secondary causes [4]. Not only do osteocytes have key functions in bone, they have also been shown to be involved in a large range of other major functions throughout the body [5], including heart, muscle and liver function, and suppressing breast cancer growth and metastasis in bone [6]. This highlights the critical role of the osteocyte in human health, and the importance of better understanding osteocyte signalling factors for the development of therapeutics to treat orthopaedic and systemic diseases.

A prime example of osteocyte sensing and coordination of bone physiology is in mechanoadaptation, with mechanical loading leading to enhanced bone formation and unloading leading to bone loss [7]. In response to macroscale deformation of bone, resident osteocytes sense the micro-mechanical environment consisting of oscillatory fluid flow-induced shear stress and relay this biophysical signal to effector cells [8]. Mechanically-stimulated osteocytes can enhance the bone forming capacity of osteoblasts via direct cell-cell contact [9], in addition to secreted factors as demonstrated by conditioned media experiments [10, 11]. Furthermore, this same mechanically-activated osteocyte conditioned media was also shown to inhibit osteoclast formation [12, 13]. Due to the non-proliferative state and short lifespan of mature bone cells, continuous bone formation requires the replenishment of the exhausted osteoblast from a mesenchymal stem/stromal cell (MSC) population [14]. Interestingly the osteocyte has also been shown to coordinate MSC behaviour, with conditioned media from mechanically stimulated osteocytes enhancing MSC proliferation, recruitment and osteogenic differentiation, demonstrating the far reaching influence of this cell type, particularly in response to a mechanical stimulus [10].

The means by which osteocytes coordinate this mechanoadaptation of bone is of great interest, with several key factors identified as playing a role in this regard and, therefore, targeted as therapeutics. There has been a plethora of studies investigating various osteocyte derived factors released in response to fluid shear, including nitric oxide (NO), prostaglandin E_2_, ATP, RANKL, osteoprotegerin (OPG) and macrophage colony-stimulating factor (M-CSF) [1]. One factor that has gained much interest is sclerostin (SOST) which is released by osteocytes and inhibits Wnt-mediated bone formation. SOST expression is inhibited following mechanical loading and inhibition of this protein via anti-sclerostin therapy has been shown in clinical trials to increase bone mineral density and reduce fracture risk [15]. To gain a greater understanding of the factors expressed by physically stimulated osteocytes, others have taken a more global approach, utilising microarrays to study global gene expression in osteocytes subjected to cyclic compressive forces [16] and osteocytes isolated from murine trabecular bone following vertebrae loading [17]. Furthermore, a proteomic analysis has been combined with a transcriptomic analysis of osteocytes subjected to fluid shear to investigate protein as well as gene expression information and reveal novel interactions between them [18]. These studies revealed the altered proteome of the osteocyte, due to fluid flow stimulation, and identified a range of proteins which may be involved in mechanotransduction, including nucleoside diphosphate kinase and calcyclin, which are of interest due to their roles in ATP and calcium-binding, respectively. However, to date, the full secretome protein signature of the osteocyte and how this is altered in response to mechanical stimulation is unknown.

A route of cell-cell communication which has garnered much attention of late is via extracellular vesicles (EVs). EVs are spherical proteolipids bi-layer surrounded vesicles secreted from cells and are involved in cell-cell communication. EVs can apparently transfer cargo including lipids, proteins and nucleic acid from one cell to another, thereby influencing the recipient cell function [19]. Interestingly, it has recently been shown that bone cells release EVs and utilise these vesicles as a mechanism to mediate osteoblast and stem/stromal cell osteogenesis [20–23]. Moreover, osteocyte-derived EVs contain miRNAs known to mediate osteoblast function, highlighting a potential non-protein based role in bone cell communication [24, 25]. Bone derived EVs may also be exploited as a potential therapy for various diseases, as well as having potential for treatment of critical size bone defects [26]. Osteoblast-derived EVs loaded with bisphosphonates have been shown to inhibit osteoclast activity *in vitro* and *in vivo* [27], supporting their potential as a powerful drug delivery method. Interestingly, the release of EVs - and thus their content-may also be altered by mechanical loading. In fact, EV release into plasma increases following exercise, with a differential protein cargo in EVs from subjects after exercise compared to those at rest [28]. Therefore, a potential mechanism of osteocyte-mediated mechanoadaptation in bone may be facilitated by mechanically-activated extracellular vesicles (MAEVs).

While changes in several factors in and released by osteocytes have been shown via proteomics analysis, the specific composition and factors implicated in mechanically-mediated osteocyte paracrine signalling are yet to be elucidated. Thus, the aim of this study was to further investigate the means by which osteocytes mediate bone mechanoadaptation, with this being achieved by constructing, for the first time, an extensive map of the osteocyte secretome protein signature. Thus, we first validated the ability of the osteocyte secretome to induce a chemotactic and osteogenic response in hMSCs using a parallel plate flow chamber approach to mechanically stimulate osteocytes. We then conducted a proteomic analysis on the osteocyte secretome via mass spectrometry, to identify proteins released by cells under both static and dynamic culture conditions. Enrichment of gene ontology terms was investigated to elucidate the primary cellular components and processes with which the osteocyte secretome is involved, with further analysis comparing the altered protein release and most differentially expressed proteins released by mechanically-stimulated cells. This led to the discovery of mechanically-activated extracellular vesicles (MAEVs). Specifically, EVs were subsequently separated from the secretome of mechanical-activated osteocyte; characterised; and found to elicit similar trends in MSC recruitment and osteogenesis to that seen with conditioned media (i.e. whole secretome). This demonstrated a key role for osteocyte EVs in mediating hMSC behaviour, identifying a novel mechanism by which osteocytes coordinate loading-induced bone formation.

## Results

### Osteocytes regulate human MSC recruitment and osteogenesis in response to fluid shear

hMSCs were cultured in conditioned medium collected from statically (CM-S) and dynamically (CM-F) cultured osteocytes, with recruitment and osteogenic gene expression being investigated (Figure 1A). A trend of increased hMSC recruitment towards CM-S compared to control medium was observed; however, this was not significant. CM-F did, however, enhance MSC recruitment; an affect that was significantly greater than with either medium (3.2-fold; p < 0.001, n = 9) or CM-S (1.8-fold; p < 0.01, n = 9), indicating the enhanced chemotaxis displayed by MSCs towards mechanically-stimulated osteocytes. The role of osteocyte paracrine signalling in driving osteogenesis was also investigated by treating hMSCs with CM-S or CM-F for 24 h and investigating expression of osteogenic genes COX2, OCN, OPN, RUNX2 and OSX (Figure 1B). Treatment with CM-S did not significantly alter expression of any of the investigated genes in hMSCs compared to medium. hMSCs cultured in CM-F resulted in consistently increased expression of all genes evaluated, with significant fold changes of 4.6 in COX2, 5.4 in OPN and 3.4 in RUNX2 compared to medium (p < 0.001, n = 4-6). These genes were also significantly upregulated with CM-F compared to CM-S with 3.0-, 2.2- and 2.3-fold changes, respectively (P < 0.01 – 0.001, n = 4-6). There was a near-significant 3.1-fold increase in OSX with CM-F compared to medium (p = 0.07, n = 4-6), in addition to a 2.5-fold increase in OCN expression in CM-F compared to CM-S. In summary, CM-S elicits marginal increases in hMSC osteogenesis, with significant increases following CM-F treatment, supporting the importance of mechanical-loading in mediating osteocyte-MSC mechanosignaling.

**Figure 1.**
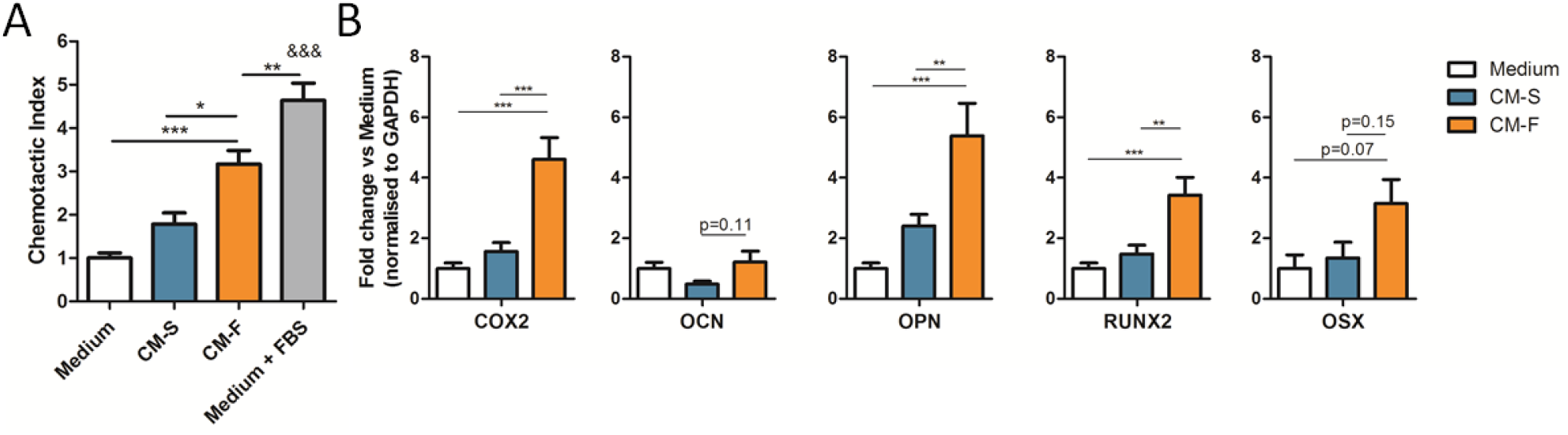
Role of osteocyte conditioned medium on hMSC osteogenesis. Migration of hSSCs towards osteocyte conditioned medium and normalised to Medium, showing significant increases in chemotactic index towards CM-F medium when compared to CM-S (n=9) (A). qPCR analysis of COX-2, OCN, OPN, RUNX2 and OSX expression in hMSCs treated with osteocyte medium from CM-S and CM-F (n=4-6) (B). Statistical analysis using using one-way ANOVA and Bonferroni’s multiple comparison post-test for chemotactic index (*p<0.05, **p□<□0.01, ***p□<□0.001, &&& p < 0.001 vs Medium and EV-S).

### Overview of identified proteins within the osteocyte secretome

Analysis of the osteocyte secretome revealed a total of 393 proteins across all groups. Within these groups, over 300 proteins were identified in both the CM-S and CM-F groups, with 112 being identified in Medium control (Figure 2C). Pearson correlations, comparing all biological replicates to one another, show that there is a high average correlation between replicates in the CM-S (0.92) and CM-F (0.90) groups (Figure 2D). When comparing CM-S and CM-F to one another, an average correlation of 0.90 is seen, revealing a significant degree of similarity in protein expression between osteocytes cultured in static and dynamic conditions. In contrast, when comparing CM groups (CM-S and CM-F) to Medium, an average correlation of 0.33 was seen between them, demonstrating the difference in the osteocyte secretome and osteocyte culture media indicating the release of proteins into culture medium from the osteocyte.

**Figure 2.**
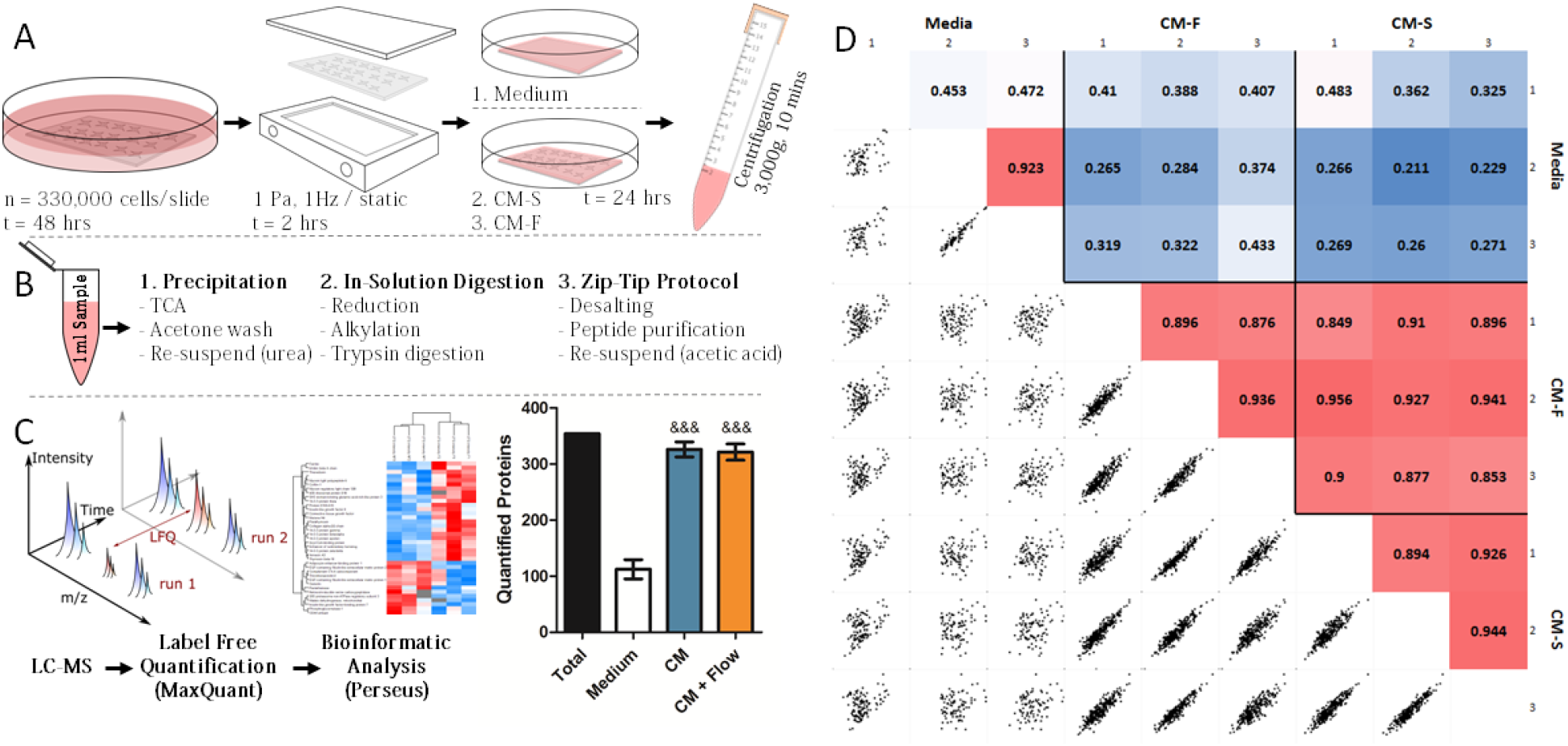
Outline of experiment procedure. MLO-Y4 cells were seeded to collagen coated glass slides and cultured for 48 hours (A), before being transferred to parallel plate flow chambers for dynamic (OFF, 1Pa, 1Hz, 2hrs) or static culture. The slides were then transferred to culture dishes and 2.5ml of serum free medium was applied, with a control group being present with collagen coated glass slides without cells. The serum free medium was collected and centrifuged to remove debris. 1ml of each sample was collected, and proteins were precipitated and digested in solution before being purified via C18 stage tips (B). Samples were analysed via LC-MS, and label free quantification was carried out in MaxQuant before a bioinformatic analysis was completed in Perseus (&&& p < 0.001 vs Medium using one-way ANOVA and Bonferroni’s multiple comparison post-test)(C). Pearson correlations between technical replicates, biological replicates and sample groups were determined, with correlations between biological replicates with combined technical replicates shown (D).

### Proteomic analysis of the osteocyte secretome reveals enrichment of proteins associated with EVs

Hierarchical clustering revealed three primary groups of protein expression within the samples. CM-S and CM-F groups comprise one of the main clusters (Figure 3A), where it can be seen that there is considerable similarity of protein content in terms of LFQ intensity within these groups. Medium samples comprise the remaining column clusters, where the reduced number and expression of proteins are more apparent when considering data without imputation (Figure 3B). Due to the similarity between osteocyte conditioned medium groups, also verified via PCA (Figure S 2), an analysis was first undertaken by combining CM-S and CM-F (termed CM), and comparing it to Medium to identified the proteins which comprise the osteocyte secretome. The results of this reveal the presence of 97 proteins which have significant differential expression in CM, indicated in red in Figure 3C, and listed in Table 1. Within these proteins, significant enrichment (enrichment factor > 1.7, p < 10^-4^) of several “extracellular” GOCC terms was shown in comparison to the total 393 identified proteins using Fisher’s exact test, with enrichment of UniProt keywords “secreted” and “signal” (enrichment factor > 1.6, p < 10^-5^) also occurred (Figure 3D). This validates the successful isolation of proteins released by the osteocyte into their surrounding environment, with evidence for further downstream signalling functions. Functional enrichment within CM proteins of GOCC terms with reference to the whole *Mus musculus* genome further reported the significant enrichment of membrane-bound vesicles and exosomes in the secretome (Table S 2). This suggested a potential role for EVs, and in particular exosomes (FDR < 10^-40^), in transporting signalling factors released by osteocytes. Functional enrichment of GOBP, GOMF and Pfam terms was also investigated, showing significant roles for these proteins in mechanosensensing and mechanosignaling, as evidenced by the most significantly enriched terms “response to stress” (FDR < 10^-6^) and “protein complex binding” (FDR < 10^-8^). The interaction network between identified proteins in the osteocyte secretome reveals a highly significant degree of protein-protein interaction (p < 10^-16^) as illustrated in Figure S 1. Enrichment analyses was also conducted on proteins more abundantly expressed in the control Medium samples using Fisher’s exact test (Figure S 3) and functional enrichment (Table S 4), revealing enrichment of muscle and cytoskeletal terms. These associations are likely due to the incorporation of proteins from rat tail collagen type 1 used for coating glass slides.

**Figure 3.**
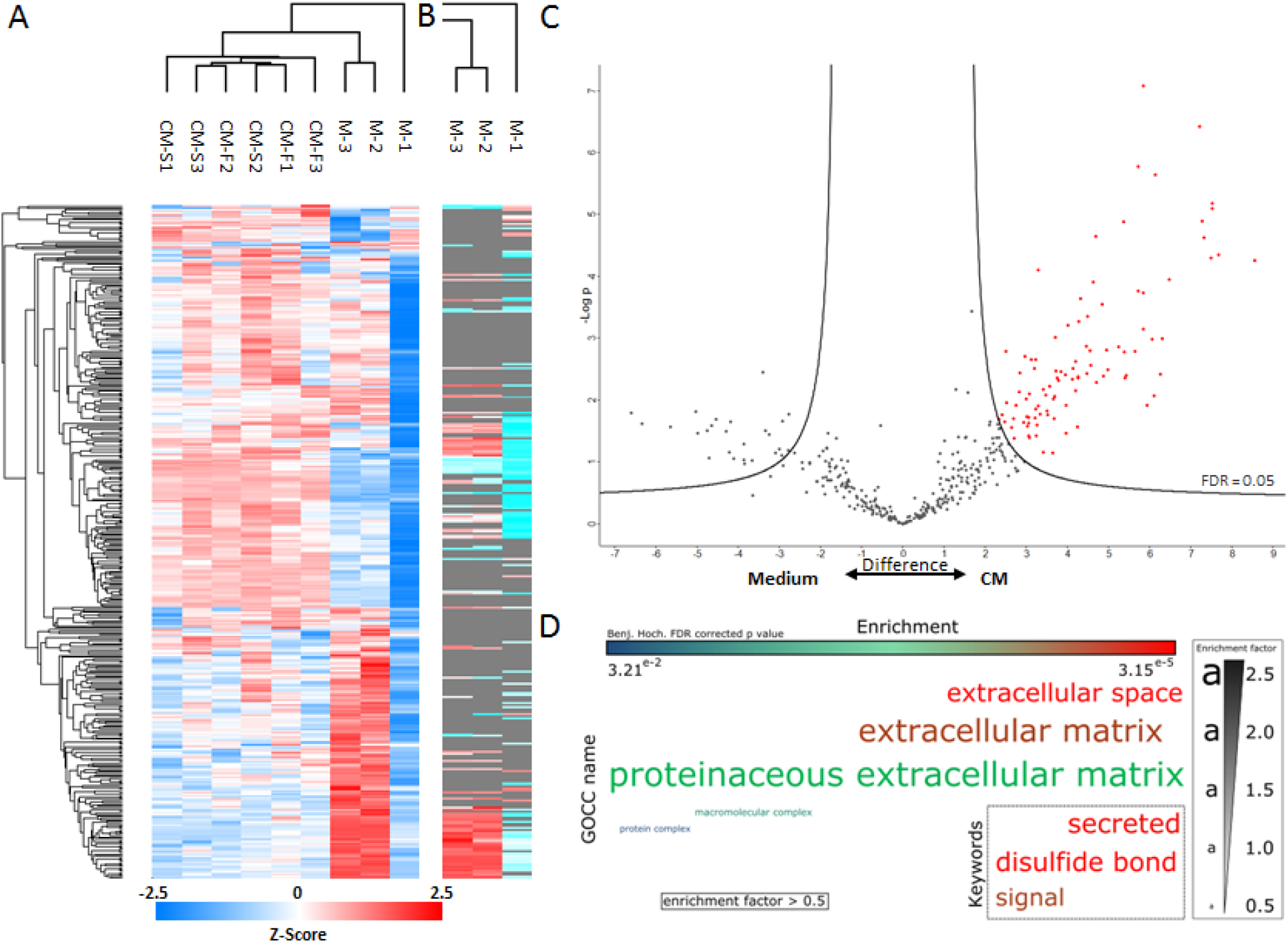
Proteomic analysis of the osteocyte secretome. Hierarchical clustering of all samples with imputed data (A) and hierarchical clustering in the control samples without imputation of data (B). Volcano plot illustrating proteins significantly upregulated proteins marked in red in CM-S and CM-F groups compared to the control (C). Enrichment analysis of GOCC terms and Uniprot keywords in upregulated proteins using Fisher’s exact test represented as a word cloud (D). The size of the word represents enrichment of terms, while colour represents FDR corrected p value. All terms with a minimum of 0.5 enrichment factor and 0.05 FDR corrected p value were included.

**Table 1.**
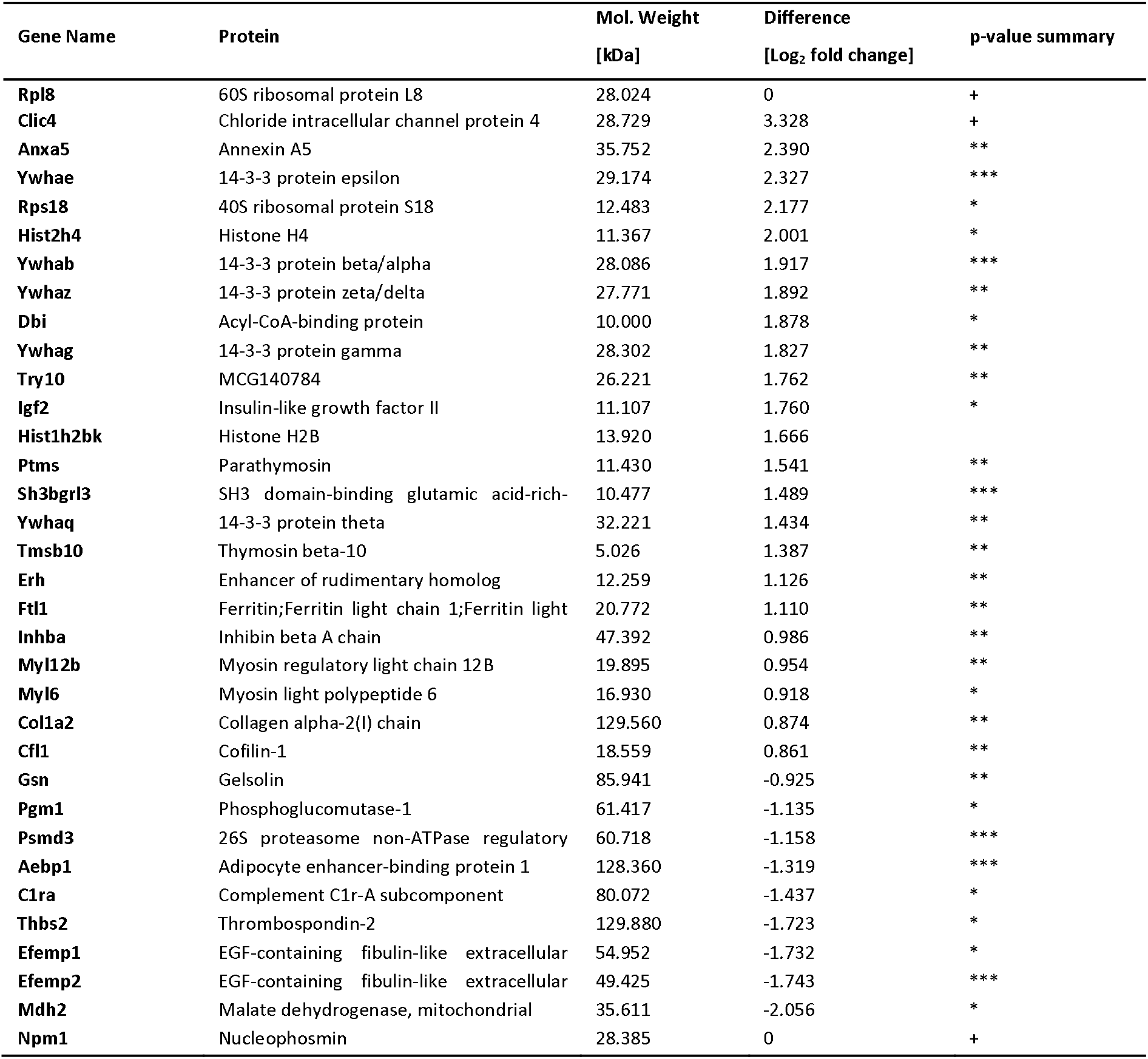
Differentially expressed proteins between CM-S and CM-F groups. *p<0.1, **p<0.05, ***p<0.01 and + indicates proteins with only a single or no detection in CM-S group (top of table) and no detection in CM-F group (bottom of table) where p-value cannot be defined.

### Mechanical stimulation alters the protein release characteristics in osteocytes

Subsequent analysis separating the CM-S and CM-F groups showed that different proteins were released from statically-cultured and mechanically-stimulated osteocytes, highlighting the role of external mechanical forces in regulating the osteocyte secretome. The more stringent criteria of only considering proteins identified in all three biological replicates in at least one of the groups reduced the total number of proteins of interest to 317. A total of 34 proteins were identified with varying degrees of significance and differential expression between groups, with 32 of these indicated on a volcano plot (Figure 4A), and a further 2 not present on the plot due to being present in only one of the CM groups. LFQ intensities of some of the most differentially expressed proteins with greater expression in CM-S (Figure 4B-D) or CM-F (Figure 4E-F) are highlighted. Of note is the enrichment of 14-3-3 proteins, all of which are upregulated in CM-F (log2 fold change = 1.43 – 2.33). Of particular interest are annexin A5 (log2 fold change = 2.39), which is associated with EVs and blood microparticles suggesting a role in systemic signalling, and histone H4 (log2 fold change = 2.00) which is associated with osteogenic growth peptide (OGP) and known to stimulate osteoblast activity [29].

**Figure 4.**
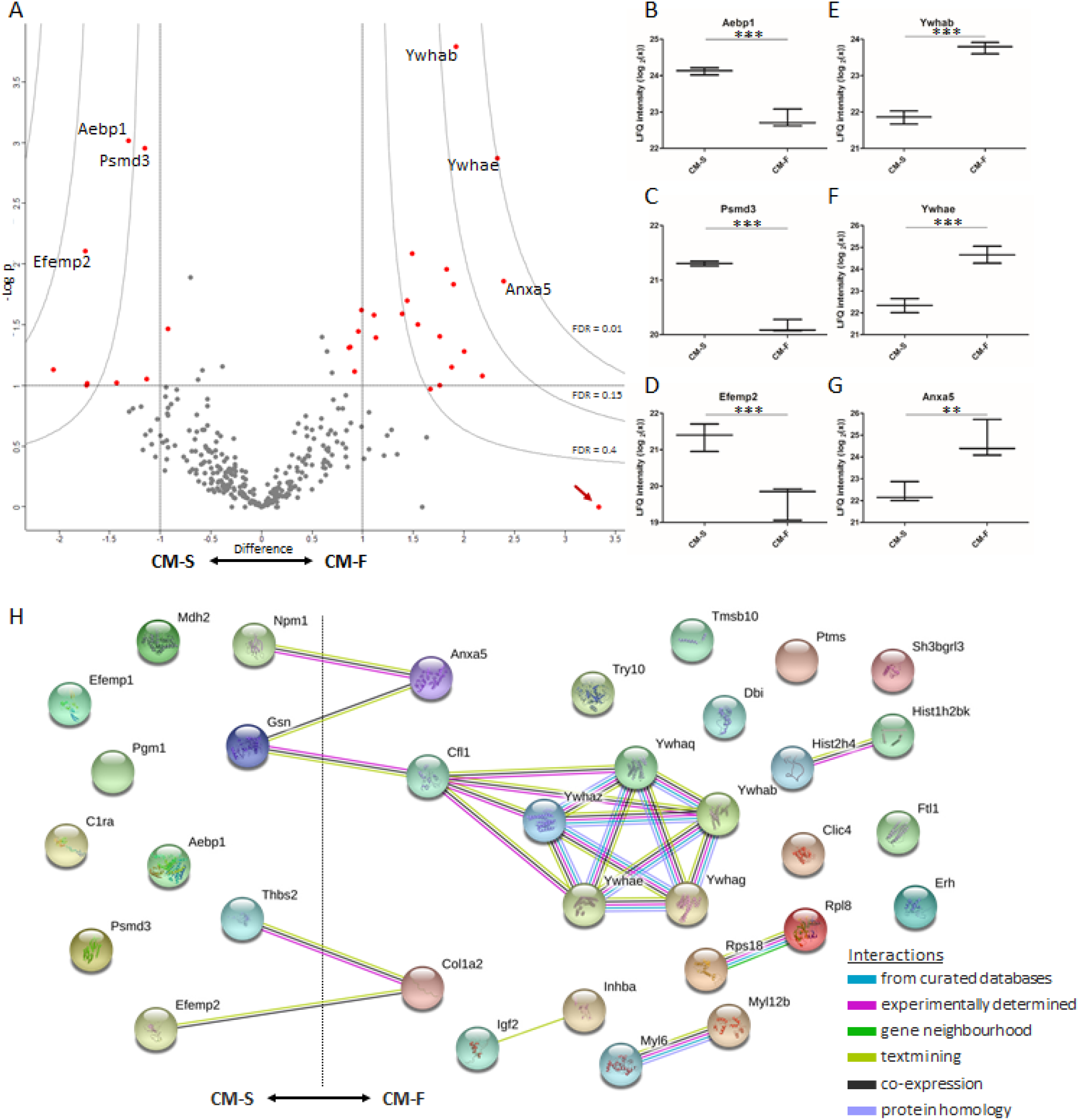
Quantifying differences between the static and dynamic osteocyte secretome. Volcano plot (A), illustrating upregulation with flow to the right and downregulation to the left. The y-axis displays the −log10 of p-value, where the horizontal line corresponding to a p-value of 0.1. Vertical lines indicate a log2 fold change of ±1. Curves illustrate indicated FDR values with S0 parameter set to 2. Whisker plots of three significantly upregulated proteins in the presence of fluid flow are indicated (B-D). The arrow indicates a protein which displays low significance due to being present in only one of the CM replicates. String DB network illustrating interactions between mechanically regulated proteins, with significant degree of protein-protein interaction (p < 10-3) (H).

Subsequently, functional enrichment in differentially secreted proteins between CM-F and CM-S was investigated to help further elucidate their collective biological relevance in mechanically mediated osteocyte signalling (Table S 3). The top four enriched GOCC terms: extracellular region, membrane-bounded vesicle, extracellular region part and extracellular exosome are associated with EV proteins with a highly significant false discovery rate (FDR < 10^-10^). 65 – 76% of all differentially secreted proteins were associated with these terms. This confirms that EVs are not only implicated in the osteocyte secretome, as demonstrated above, but are a key component of mechanically-mediated signalling. Also of substantial interest is the enrichment of the top two GOMF terms “calcium ion binding” (FDR < 0.01) and “phosphoserine binding” (FDR < 0.05), revealing the potential role of mechanically-activated osteocyte EVs as sites of mineralisation via binding of calcium and phosphate components. A String DB network was constructed to further investigate any potential interactions between proteins associated with EVs (Figure 4H) revealing a significant degree of protein-protein interaction (p < 10^-3^). Interestingly, there are several interactions between positively and negatively regulated proteins, including an interaction path between Anxa5 and Ywhab/Ywhae which are associated with calcium ion binding and phosphoserine binding, respectively. Between these nodes are gelsolin and cofilin, the former of which is calcium sensitive and both of which have been shown to regulate changes in the actin cytoskeleton [30], as well as occurring in vesicles from mineralising osteoblasts [31].

### EVs are present within the osteocyte secretome and EV morphology and size distribution is not altered by mechanical stimulation

Given the identification of EV-associated proteins within the osteocyte secretome, we next investigated whether osteocytes release EVs and, if so, whether EV characteristics were altered by mechanical stimulation. EVs were successfully separated from osteocyte CM using filtration and ultracentrifugation, with the presence of EVs confirmed by TEM imaging and immunoblotting. TEM imaging confirmed the presence of EVs of typical morphology and size (Figure 5A-B). The presence of EVs was further confirmed via immunoblotting, with no detection of negative marker GRP-94, and detection of positive markers TSG101 and ALIX (Figure 5C). EV concentration was not significantly different between EVs separated from the CM-S (EV-S) and EVs separated from the CM-F (EV-F), both being within the range of 0.8 – 2.6 μg/ml, and with average values of 1.2 μg/ml and 1.5 μg/ml, respectively (Figure 5D). It can be seen that there is a change in particle size distributions between EV-S and EV-F (Figure 5E), however, no changes in average particle size was detected, with values of 177 nm and 183 nm respectively (Figure 5F).

**Figure 5.**
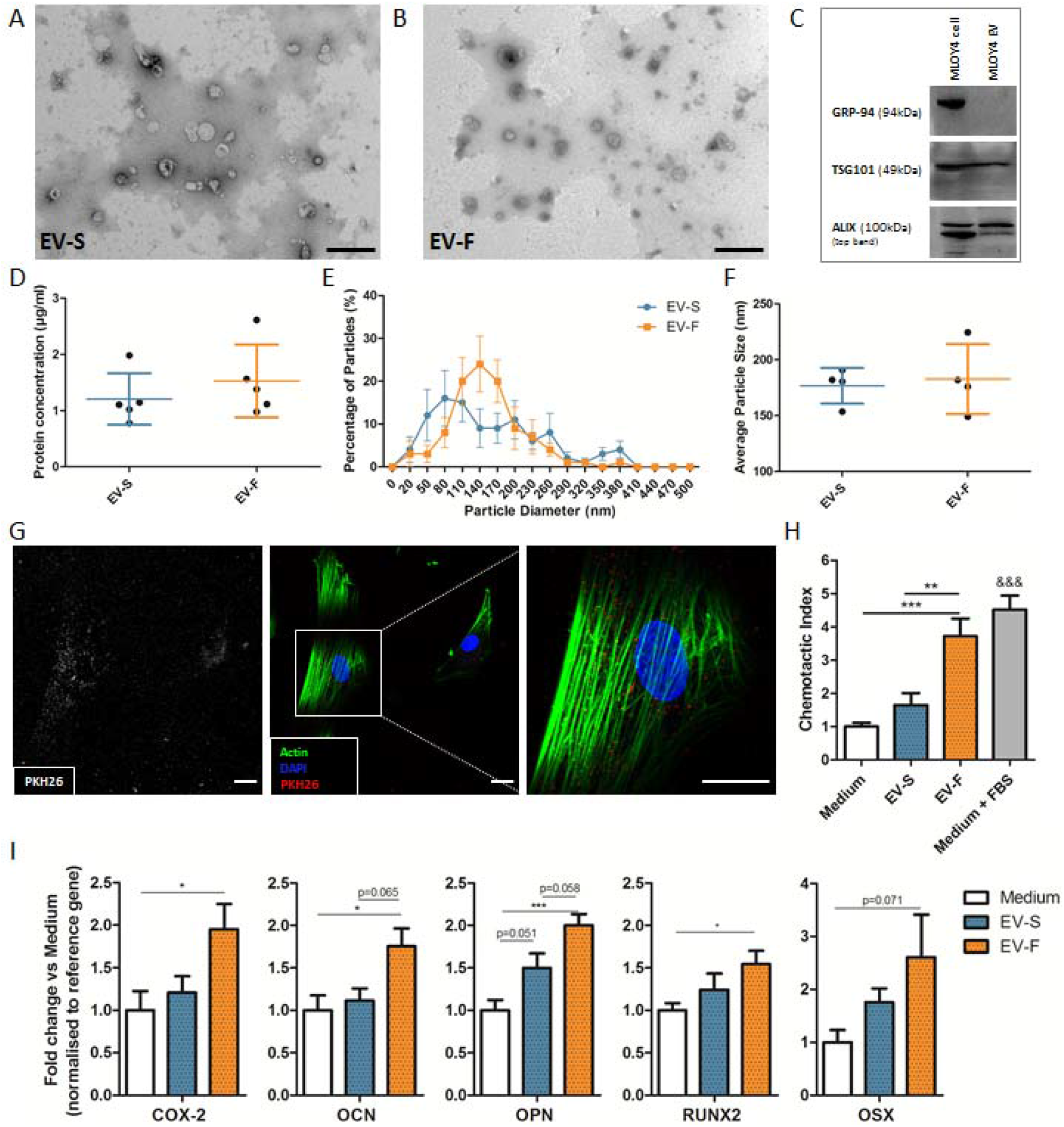
Characterisation of EVs and their influence on hMSC osteogenesis. TEM image of EVs isolated from osteocyte CM-S (A) and CM-F (B). Immunoblots confirmed the presence of EVs via negative marker GRP-94 and positive markers TSG101 and ALIX (C). Protein concentration of EVs in conditioned medium groups (n=5) (D). Nanoparticle size analysis on EVs confirmed no significant difference in distribution (E) or average size (F) between groups (n=4). Immunofluorescent images illustrating osteocyte EV uptake by hSSCs, as demonstrated by localisation of PKH26 labelled EVs within the cell body (Scale = 10μm) (G). Migration of hSSCs towards EVs isolated from osteocyte conditioned medium and normalised to Medium, showing significant increases in chemotactic index towards CM-F medium when compared to CM-S (H). qPCR analysis of COX-2, OCN, OPN, RUNX2 and OSX expression in hSSCs treated with EVs from osteocyte medium from CM-S and CM-F (I). Statistical analysis using using one-way ANOVA and Bonferroni’s multiple comparison post-test (*p<0.05, **p□<□0.01, ***p□<□0.001, &&& p < 0.001 vs Medium and EV-S)

### Osteocytes regulate human MSC recruitment and osteogenesis in response to fluid flow shear via MAEVs

To determine whether mouse osteocyte-derived EVs could be taken up by hMSCs, we labelled EVs with PKH26. Following 24 hr treatment, labelled-EVs were preferentially located within the cytoplasm, indicating uptake of EVs by hMSCs (Figure 5G). Control samples are illustrated in Figure S 4. A high density of EVs can be seen around the nuclear region in particular with minimal detection within the nucleus.

Upon verifying EV uptake, the cellular response of hMSCs subjected to EVs separated from CM-S (i.e. EV-s) or CM-F (EV-F) was investigated. Specifically, hMSCs were treated with EV-S and EV-F to investigate recruitment and osteogenic gene expression as previously demonstrated with CM. EV-S resulted in a slight non-significant increase in MSC recruitment (Figure 5H), while this response was significantly enhanced with EV-F, yielding a 3.7-fold increase compared to medium (p < 0.001, n = 9) and 2.3 fold increase compared to EV-S (p < 0.01, n = 9). This trend closely mirrored that seen with whole secretome. Osteogenic gene expression (Figure 5I) showed a consistent trend of marginally increased expression with CM-S treatment, which was further enhanced with CM-F. There was a near-significant increase of 1.5-fold in OPN (p = 0.051, n = 17-18) when comparing CM-S to Medium. CM-F resulted in significant changes compared to medium of 2.0-fold in COX2 (p < 0.05, n = 17-18), 1.8-fold in OCN (p < 0.05, n = 14-15), 2.0-fold in OPN (p < 0.001, n = 16-18) and 1.5-fold in RUNX2 (p <0. 05, n = 20), with a near-significant increase of 2.6-fold in OSX (p = 0.07, n = 21-23). In addition, near-significant increases in OCN and OPN were detected comparing EV-F and EV-S. Moreover, orthology searches using the basic local alignment search tool BLAST [32] were performed for the human gene sequences (COX2, OCN, OPN, RUNX2, OSX) in the murine genome (version: *Mus musculus* GRCm38.p4). The bioinformatic tool, *in silico* PCR (UCSC Genome Browser), was used to confirm the lack of amplification of the human primer sequences in the murine genome [33], confirming that amplified genes are human, and not due to possible transfer of murine mRNA from the MLO-Y4 cell line. In summary, there is a trend of increasing osteogenesis in hMSC following EV-S treatment. However, this affect becomes significantly greater with EV-F treatment, showing a similar trend to that seen with whole secretome and demonstrates that EVs from mechanically-activated osteocytes are key drivers of stem/stromal cell recruitment and osteogenesis.

## Discussion

Osteocytes are mechanosensitive cells which play a fundamental role in coordinating loading-induced bone formation via the secretion of paracrine factors which drive effector cell behaviour. One of the most important of which are bone marrow mesenchymal stem/stromal cells (MSCs) which are responsible for replenishing the bone forming osteoblast population. However, the exact mechanisms by which osteocytes relay mechanical signals to these cells are poorly understood. A greater understanding of these mechanisms would thus have profound implications for the development of therapies to treat the wide range of diseases with which the osteocyte has been linked, one of the most devastating of which is osteoporosis. Therefore, this study aimed to demonstrate the potency of the mechanically stimulated osteocyte secretome in driving human MSC behaviour, and fully characterise its contents with the aim of identifying the key secreted factors regulating bone mechanobiology. Herein, we demonstrate that osteocytes subjected to oscillatory fluid shear secrete factors that significantly enhance hMSC recruitment and osteogenesis. To uncover the osteocyte derived secreted factors which drive hMSC behaviour, we performed a proteomic analysis of the osteocyte secretome to uncover an extensive map of proteins which are released both under static conditions and following mechanical stimulation. Over 300 proteins comprising the osteocyte secretome were identified with 34 proteins differentially expressed following mechanical stimulation. The osteocyte secretome was significantly enriched with proteins associated with extracellular vesicles (EVs) and exosomes indicating a role for secreted vesicles in mediated mechanically driven osteocyte-MSC communication. EVs were subsequently separated from the mechanical activated osteocyte secretome, characterised, and found to elicit similar trends in MSC recruitment and osteogenesis to that seen with conditioned media, demonstrating a key role for osteocyte EVs in mediating hMSC behaviour.

Mechanically stimulated osteocytes secrete paracrine factors that recruit human MSCs and enhance osteogenesis. The ability of mechanically stimulated osteocytes to influence MSC behaviour is in agreement with previous findings *in vivo* where mechanical loading of bone results in the recruitment and osteogenic differentiation of endogenous [34] or transplanted exogenous osteoprogenitors [35]. Furthermore a similar trend in recruitment has been shown in murine MSCs where a 128% increase in recruitment was observed following exposure to conditioned media collected from osteocytes cultured on a rocking platform [10]. Interestingly in the same study, mechanically activated osteocyte conditioned media was also been shown to induce osteogenesis of MSCs as demonstrated by upregulation of Opn and Cox-2 gene expression and enhanced mineral deposition [10, 11]. We have demonstrated a comparable increase in COX-2 and OPN expression in human MSCs, as well as increases in OCN, OSX and RUNX2. These findings, along with other previous work investigating the effect of the osteocyte secretome on osteoblast proliferation, migration and osteogenesis [9, 36], further reinforce the importance of the osteocyte secretome and its contents in the indirect biophysical regulation of MSCs and loading-induced bone formation [37, 38].

To determine the mechanisms by which osteocytes coordinate MSC recruitment and osteogenesis in response to loading, for the first time, we identified a detailed a map of the osteocyte secretome via a mass spectrometry based proteomic analysis. Interestingly, a number of key proteins, such as Sclerostin, which is known to be secreted by osteocytes, were not detected. This may due to the limitation of the MLO-Y4 osteocyte cell line but given the pro-osteogenic effect of this mechanically activated osteocyte secretome, this opens the possibility of identifying novel factors regulating MSC behaviour. We have identified several proteins from a previous proteomic analysis on osteocyte lysates [18], revealing potential roles for these proteins in cell signalling. Further analysis sought to investigate the role of mechanical forces on the contents of the osteocyte secretome, with differential expression of a range of proteins being identified compared to statically cultured cells. One Pfam group of particular interest which are significantly enriched with fluid shear are the 14-3-3 proteins. One study reports that 14-3-3 beta has a negative effect on osteogenesis, with downregulation in calvaria organ cultures resulting in increased bone formation [39]. 14-3-3 epsilon is released by osteoblasts/osteocytes in response to dynamic compression, inducing the release of catabolic factors in chondrocytes in a dose dependent manner, mimicking the effect of compression [40]. Interestingly, TAZ, a known mechanosensor and transcriptional modulator, has also been linked to 14-3-3 proteins, with decreased binding being shown to result in increased TAZ nuclear localisation [41], further indicating a role for 14-3-3 proteins in mechanically mediated signalling in bone. Other proteins of particular interest are histone H4 and annexin A5. The acetylation of histone H4 has been shown to promote the induction of osteocalcin gene expression in osteoblasts [42], with histone deacetylase inhibition being shown to promote osteoblast differentiation [43] and increase mineralisation [44]. More specifically, the C-terminus of histone H4, termed osteogenic growth peptide (OGP), a circulating stimulator of osteoblast activity [29], plays key roles in regulating the behaviour of bone residing cells, such as stimulating proliferation, phosphatase activity and mineralisation of osteoblasts and proliferation and osteogenic differentiation of MSCs [45]. Annexin A5 has been shown to increase at the cell membrane in osteoblasts under fluid flow, with Ca^2^+ ion levels also being seen to increase. It was found that the disruption of annexin A5 inhibited Ca^2+^ levels, implicating its role in calcium signalling [46], with its knockdown in osteoblasts impairing proliferation, ALP expression and Runx2 expression [47]. One downregulated protein of interest is thrombospondin 2. The knockdown of this protein in mice increases angiogenesis [48] with endosteal bone formation being shown in another study to increase as a result of increased bone marrow derived osteoprogenitors [49]. Thrombospondin 2 null mice have also demonstrated enhanced callus bone formation, vascularity and MSC proliferation following tibial fracture [50]. We have identified a list of proteins released by the osteocyte, many of which are mechanically regulated, and have been linked to bone physiology. This therefore represents a database of proteins to help better understand the osteocyte coordination of bone anabolism and catabolism and provides a list of potential therapeutic targets to mimic this behaviour.

Functional enrichment analysis of the osteocyte secretome revealed a strong association with ‘extracellular exosome’ and ‘membrane bound vesicles’ with 66% of secreted proteins being linked to these cellular organelles. Moreover, in response to mechanical stimulation, 70% of the mechanically regulated proteins were associated with extracellular vesicles. EVs were successfully separated from osteocyte conditioned media and interestingly we did not detect any changes in EV morphology or quantity between static and dynamic groups, in contrast with previous work which has demonstrated an upregulation in EV number following fluid shear stimulation [51]. We have however demonstrated an almost identical trend in MSC recruitment with CM and EVs, both of which are enhanced following fluid shear, providing evidence for the key role of osteocyte derived EVs in mediating this response. Similarly, the almost identical trends in MSC osteogenic gene expression treated with CM and EVs further provide evidence for the role of osteocyte derived MAEVs in facilitating cell-cell communication in bone. Given the similar concentrations of EVs between groups, it is expected that this pro-osteogenic effect is a result of EV content changing in response to mechanical stimulation. Many of the proteins identified in this paper have also been identified in a proteomic analysis of osteoblast released EVs [31, 52]. A previous *in vivo* study has suggested a role for EVs in systemic signalling, demonstrating altered miRNA expression in EVs isolated from the plasma of osteocyte ablated mice and wild-type mice [24], while other studies have demonstrated the potential for EVs in therapeutics to enhance osteogenic gene expression [53], the use of drug loaded EVs for osteoporosis therapies [27], and the use of EVs for functionalisation of TE scaffolds to enhance bone regeneration [54, 55]. In addition to osteocyte derived MAEVs and the contents driving MSC osteogenesis, MAEVs may also act as sites for mineral nucleation, as has previously been demonstrated in osteoblast EVs [56, 57]. Of particular interest in this regard is the enrichment of calcium ion binding (such as annexin A5) and phosphoserine binding (such as 14-3-3 proteins Ywhae and Ywhab) proteins in osteocyte MAEVs, which we have shown to be linked in a protein interaction network. Annexin A5 is linked to the calcium sensitive protein gelsolin [30], which in turn is linked to the 14-3-3 proteins via the phosphate regulating cofilin [58], both of which regulate changes in the actin cytoskeleton [30]. In addition to the known role of calcium ions in mineralisation, negatively charged amino acids such as phosphoserine are also known to play a key role in hydroxyapatite nucleation and growth [59]. Therefore osteocyte EVs may promote mineralisation via delivery of calcium and phosphate interacting proteins through interaction with gelsolin and cofilin respectively. Taken together, we have identified mechanically activated extracellular vesicles as a key mechanism by which osteocyte communicate chemotactic and osteogenic signals to osteoprogenitors in response to loading, highlighting these osteocyte derived MAEVs as a potential cell free therapy to mimic the beneficial effect of loading and enhance bone formation.

One of the limitations of this study is the use of the MLO-Y4 cell line. While these cells largely behave as osteocytes in vivo, releasing various characteristic signalling molecules and mediating bone cell behaviour [60, 61], they lack several characteristics such as the typical absence of sclerostin expression and low DMP-1 expression [62]. In spite of these limitations, we have demonstrated the potent effect this cell can have on hMSC osteogenesis and have identified novel factors which are released to achieve this. These cells are also well characterised in the literature, allowing direct comparison of results with a wide range of studies.

In summary this study presents evidence that the mechanically stimulated osteocyte secretes factors which coordinates MSC recruitment and osteogenesis demonstrating a mechanism required for loading-induced bone formation. Importantly, for the first time, we have mapped the osteocyte protein secretome and determined how this is altered in response to mechanical stimulation generating a database of potential factors mediating this mechanism. Lastly, this study also demonstrates the presence and fundamental role of mechanically activated EVs (MAEVs) released by osteocytes in coordinating MSC recruitment and osteogenesis, identifying a novel mechanism by which osteocytes coordinate bone mechanobiology. Moreover, these pro-osteogenic osteocyte derived MAEVs represent a potential cell-free therapy to enhance bone regeneration and repair in diseases such as osteoporosis.

## Materials and Methods

### Cell culture

MLO-Y4 osteocyte like cells (Kerafast) [63] were cultured, as previously described [64], in α-MEM growth medium with 2.5% fetal bovine serum (FBS), 2.5% calf serum (CS), 1% penicillin/streptomycin (PS) and 1% L-glutamine during static culture and fluid shear stimulation. For whole secretome/conditioned medium CM) studies, cells were cultured in α-MEM with 1% PS and 1% L-glutamine. hMSCs were isolated from bone marrow (Lonza), characterised by tri-lineage differentiation (data not shown), and maintained in Dulbecco’s Modified Eagle Medium (DMEM) with 10% FBS and 1% PS unless otherwise stated. All cells were cultured at 37°C and 5% CO_2_.

### Mechanical stimulation and conditioned medium collection

48 h prior to fluid shear application, 75 x 38 mm glass slides were coated with 0.15mg/ml type I collagen (Sigma C3867) for one hour and washed with PBS, after which osteocytes were seeded at a density of 1.16 x 10^4^ cells/cm^2^. Glass slides were transferred to custom made parallel plate flow chambers (PPFC) as previously described [65]. Each glass slide was assembled within an individual PPFC under sterile conditions and incubated at 37°C and 5% CO_2_. Cells in PPFCs were either subjected to a fluid shear stress of 1 Pa at a frequency of 1 Hz, or maintained in the PPFC under static conditions, with each condition completed in quadruplicate. After two hours of treatment, slides were transferred to culture dishes, washed with PBS, and 2.5 ml of serum-free medium was applied. A control group consisting of collagen-coated glass slides with no cells was also incubated with 2.5 ml of serum-free medium. All culture dishes were incubated for 24 h and medium was collected from cells which had undergone fluid shear (CM-F), statically cultured cells (CM-S) and from cell-free slides with collagen coating (Medium). Samples were centrifuged at 3,000g for 10 mins at 4°C to remove debris, after which the supernatant was collected and stored at −80°C prior to use (Figure 2A).

Chemotaxis of hMSCs was assessed using Boyden chambers with a pore size of 8 μm (Merck Millipore, PIEP12R48). Cells were seeded on the upper membrane in serum free α-MEM medium at a density of 30,000 cells/cm^2^ and allowed to adhere 4 h before being transferred to the wells containing chemotactant (Medium (serum free), CM-S, CM-F, Medium + 10% FBS). Cells were then cultured for a further 18 h, fixed with 10% formalin solution and stained with haematoxylin. Light microscopy was used to determine the number of migrated cells, which was then normalised to Medium for each group.

### Effect of osteocyte conditioned media on bone marrow mesenchymal stem/stromal cell osteogenesis

hMSC cells were seeded in 6-well plates at a density of 6,500 cells/cm^2^ and cultured for 24 h. Osteocyte CM (CM-S, CM-F) was then applied and hMSCs were cultured for a further 24 h after which time cells were lysed with tri-reagent (Sigma Aldrich) and mRNA isolated as per the manufacturer’s protocol. RNA concentration was measured using a Nanodrop spectrophotometer and sample purity was checked via 260/280 and 260/230 absorbance ratios. 200 ng RNA was reverse transcribed to cDNA using a High-Capacity cDNA Reverse Transcription Kit (Applied Biosystems). Commercially available primers (Sigma Aldrich, KSPQ12012) were used to determine levels of cyclooxygenase 2 (COX2), osteocalcin (OCN), osteopontin (OPN), runt-related transcription factor 2 (RUNX2) and osterix (OSX) (Table S 1). qPCR was performed using a reaction volume of 20 μl containing 10 μl SYBR green PCR MasterMix (Invitrogen Ltd, Paisley, UK), 0.8 μl of each forward and reverse primer, and 8.4 μl DNase free water. Plates were run on an ABI 7500 Fast real-time PCR system (Life Technologies, Carlsbad, CA, USA).

### Sample preparation for MS analysis

Protein precipitation was carried out with 1ml of each sample (Medium, CM-S, CM-F) using trichloroacetic acid (TCA), and, following centrifugation at 18,500 g, the pellet re-suspended in 6M urea in 50mM ammonium bicarbonate. Samples were reduced with 5 mM dithiothreitol for 30min at 60°C and alkylated with 10mM iodoacetamide for 30min at room temperature in the dark, after which ammonium bicarbonate was added to bring the concentration of urea to 1.8M. The reduced and alkylated proteins were then digested overnight with trypsin at a ratio of 1:50 w/w trypsin to protein at 37°C and 350 rpm on a Thermomixer. Digestion was then stopped with 8.8M hydrochloric acid. Peptides were bound and desalted using C18 ZipTips (Merck Millipore) and washed with 0.1% trifluoroacetic acid (TFA) before being re-suspended in 10 μl elution solution (50% acetonitrile in 0.1% TFA). Samples were concentrated using a SpeedVac vacuum concentrator until roughly 4 μl remained, before being re-suspended in 20 μl 0.5% acetic acid (Figure 2B).

### LC MS/MS analysis

Biological samples (n=3) were run with two technical replicates on a Thermo Scientific Q Exactive mass spectrometer connected to a Dionex Ultimate 3000 (RSLCnano) chromatography system. Each sample was loaded onto a fused silica emitter (75 μm ID, pulled using a laser puller (Sutter Instruments P2000)), packed with UChrom C18 (1.8 μm) reverse phase media (nanoLCMS Solutions LCC) and was separated by an increasing acetonitrile gradient over 47/60 minutes at a flow rate of 250 nL/min. The MS was operated in positive ion mode with a capillary temperature of 320°C, and with a potential of 2300V applied to the frit. All data was acquired with the MS operating in automatic data dependent switching mode. A high resolution (70,000) MS scan (300-1600 m/z) was performed using the Q Exactive to select the 8 most intense ions prior to MS/MS analysis using high-energy collision dissociation (HCD).

### MS data analysis

Raw data from MS analysis was processed using MaxQuant software [66, 67] version 1.5.5.1 and spectra searched using the built in Andromeda search engine [68] with the Uniprot FASTA validated Mus musculus database being used as the forward database and the reverse for the decoy search being generated within the software. A minimum six amino acid length criteria was applied and the FDR for MS data analysis was set to 1% at the peptide and protein level. Cysteine carbamidomethylation was included as a fixed modification and oxidation of methionine and protein N-terminal acetylation were set as variable modifications for the peptide search. The “match between runs” algorithm was used to transfer peptide identifications between MS runs where possible to increase total number of protein hits. At least one unique or razor peptide was required per protein group for identification. Label-free quantification (LFQ) was carried out using the MaxLFQ algorithm [69] within the software, with Fast LFQ being disabled. Other settings were kept as default in the software.

### Extracellular vesicle isolation from conditioned media

Medium from statically and dynamically cultured osteocytes was collected and centrifuged at 3000 g for 10 min to remove debris. Medium was then filtered through a 0.45 μm pore filter. Medium was subsequently ultracentrifuged at 110,000 g for 75 min at 4°C, using an SW32.Ti swing bucket rotor. Collected EV pellets were washed in PBS and the ultracentrifugation process was repeated.

### Characterisation of extracellular vesicles

#### TEM imaging

EV imaging was conducted via a JEOL JEM1400 transmission electron microscope (TEM) coupled with an AMT XR80 digital acquisition system. Samples were physiosorbed to 200 mesh carbon-coated copper formvar grids and negatively stained with 1% uranyl acetate.

#### Immunoblotting

For immunoblotting, cell pellets and EVs were lysed using cell extraction buffer (Invitrogen, Carlsbad, CA, USA) supplemented with protease inhibitor cocktail (Roche, Basel, Switzerland). Protein quantification was performed using Bio-Rad protein assay (Bio-Rad, Hercules, CA, USA). Cellular and EV protein (8μg) were resolved on 10% SDS gels and transferred to PVDF membranes (BioRad). Blots were incubated at 4°C overnight with primary antibodies to GRP-94 (Cell Signalling, 1:2000 dilution), TSG101 (Abcam, 1:1000 dilution) and PDC6I/ALIX (Abcam, 1:1000 dilution). Secondary antibodies were incubated for 1hr at room temperature and developed using Immobilon Western Chemiluminescent HRP substrate (Millipore, MA, USA).

#### Quantification of EV content in conditioned medium

As a surrogate of EV quantities, protein contents were measured using a BCA protein assay kit (Thermo Scientific, 23227). BSA standards (10 μl) were added to a 96 well plate after which 200 μl of working reagent was added (50:1 ratio of reagents A & B). EV samples were diluted in CST lysis buffer (Cell Signaling Technology, 9803), vortexed, and incubated for 1 hr on ice. 10 μl of sample lysates were added to the plate and mixed with 200 μl of working reagent. The plate was incubated for 30 min at 37°C and absorbance read on a spectrophotometer at 562 nm. BCA assay results combined with the volume of the isolate were used to calculate the total quantity of protein in the EV isolates and this value was used to calculate the original concentration of EV protein in the conditioned medium.

#### Particle size analysis

Particle size analysis was performed on EV samples using a NTA NS500 system (NanoSight, Amesbury, UK). EV samples were diluted 1:50 in PBS and injected into the NTA system, which obtained 4 x 40 second videos of the particles in motion. Videos were then analysed with the NTA software to determine particle size.

### Uptake of EVs by MSCs

For fluorescent labelling, 2 μg of EVs were incubated with 2 μM PKH26 dye solution (PKH26GL, Sigma) for 5 mins, after which staining was inhibited via addition of 1% BSA solution for 1 minute. Labelled EVs were pelleted, the excess dye solution aspirated, and washed twice with culture medium. hMSCs were seeded at a density of 10,000 cells/cm^2^ in Nunc glass-bottomed dishes (150680, Thermo Fisher) and cultured for 24 h. Cells were washed with PBS before being incubated with either PKH26-labelled EVs or a dye control containing no PBS with no EVs. Cells were fixed after 18 h and stained with Alexa Fluor 488 phalloidin (1:40) (A12379, Thermo Fisher) and DAPI (1:2000) (D9542, Sigma) to label the actin cytoskeleton and nuclei before mounting with Fluoroshield (F6182, Sigma) and imaged using confocal microscopy.

### Statistical and bioinformatics analysis

Statistical analysis on recruitment and gene expression data was carried out using using one-way ANOVA and Bonferroni’s multiple comparison post-test (*p<0.05, **p□<□0.01, ***p □<□ 0.001. &&& p < 0.001 of positive control compared to all other groups).

Bioinformatic analysis was performed using Perseus 1.5.5.3 [70] to analyse LFQ data from MaxQuant. Potential contaminants, proteins identified in the decoy reverse database and proteins identified only by site modification were omitted. LFQ values were transformed using a log_2_(x) function. For clustering and principal component analysis (PCA), imputation was carried out (width = 0.3, down shift = 1.8) where missing values were replaced by values from a normal distribution. For hierarchical clustering, log transformed intensities were normalised by z-score and clustered using the Euclidean distance method for both columns and rows. Pathway enrichment analysis of clusters was carried out using a Fisher’s exact test with the Benjamini-Hochberg FDR threshold set to 5%, with gene ontology cellular component (GOCC), biological process (GOBP), molecular function (GOMF) and UniProt keywords being analysed for enrichment. A Student’s T-Test with a permutation-based FDR (1, 15, 40%) was carried out to identify differences in expression of proteins between groups, and volcano plots constructed with difference (log2 fold change) on the x-axis and significance (-log10 transformed) on the y-axis. The difference on the x-axis corresponds to the difference between the mean expression values of log2 transformed data, where a difference of n corresponds to fold change of 2^n^. Pathway enrichment analysis was carried out these significantly upregulated proteins using the Fisher’s exact test with Benjamini-Hochberg FDR cut-off of 5%. Results were represented as word clouds, with the size of the word representing degree of enrichment and colour representing FDR corrected p value. All terms with a minimum of 0.5 enrichment factor were included. StringDB [71] was used to generate protein-protein interaction networks of differentially-expressed proteins and perform functional enrichment analysis of gene ontology and protein family (Pfam) terms. For further analysis between CM-S and CM-F groups, only proteins that were identified in all three biological replicates if at least one of the groups were considered for further analysis.

## Supplementary figures and tables

**Table S 1.**
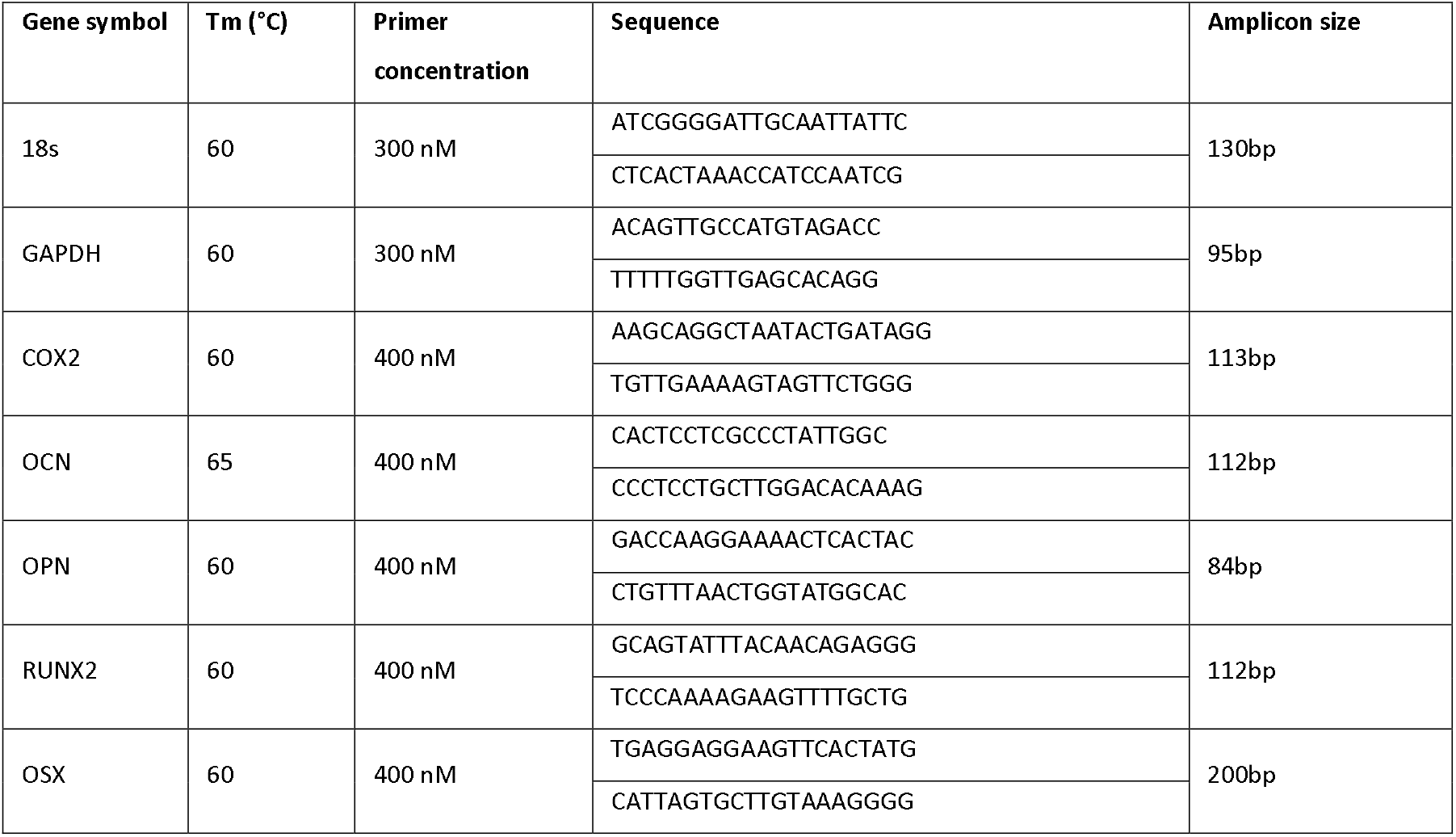
Primer sequences and concentrations employed in quantitative PCR analysis.

**Table S 2.** Functional enrichments in CM proteins indicated in Figure 3C using String DB, with observed gene count out of a total of 105 genes and FDR cut-off of 2%. (note: 105 genes were identified from 97 proteins)

| Pathway description | Observed gene count | False discovery rate (FDR) |
| --- | --- | --- |
| <u>GOCC (Gene Ontology Cellular Component)</u> |  |  |
| extracellular exosome | 71 | 1.72E-41 |
| extracellular region part | 76 | 5.05E-39 |
| extracellular region | 77 | 1.20E-35 |
| membrane-bounded vesicle | 70 | 2.07E-34 |
| extracellular space | 47 | 7.88E-32 |
| extracellular matrix | 26 | 9.57E-22 |
| proteinaceous extracellular matrix | 22 | 5.85E-18 |
| cytoplasmic membrane-bounded vesicle | 26 | 1.80E-12 |
| cytoplasmic vesicle | 24 | 9.53E-10 |
| myelin sheath | 12 | 9.48E-09 |
| <u>GOBP (Gene Ontology Biological Process)</u> |  |  |
| response to stress | 36 | 6.92E-07 |
| antigen processing and presentation of peptide antigen via MHC class I | 6 | 4.09E-06 |
| protein folding | 10 | 8.52E-06 |
| response to wounding | 13 | 2.05E-05 |
| regulation of biological process | 63 | 5.41E-05 |
| pyruvate metabolic process | 6 | 1.73E-04 |
| extracellular matrix organization | 9 | 1.73E-04 |
| biological regulation | 63 | 1.99E-04 |
| glycolytic process | 5 | 2.85E-04 |
| response to endogenous stimulus | 19 | 2.85E-04 |
| <u>GOMF (Gene Ontology Molecular Function)</u> |  |  |
| protein complex binding | 20 | 7.80E-09 |
| protein binding | 52 | 7.85E-09 |
| binding | 73 | 5.31E-08 |
| RNA binding | 26 | 3.35E-07 |
| poly(A) RNA binding | 23 | 3.35E-07 |
| isomerase activity | 9 | 9.02E-06 |
| macromolecular complex binding | 21 | 1.05E-05 |
| receptor binding | 20 | 1.22E-05 |
| calcium ion binding | 15 | 1.22E-05 |
| peptide binding | 9 | 1.90E-05 |

**Table S 3.**
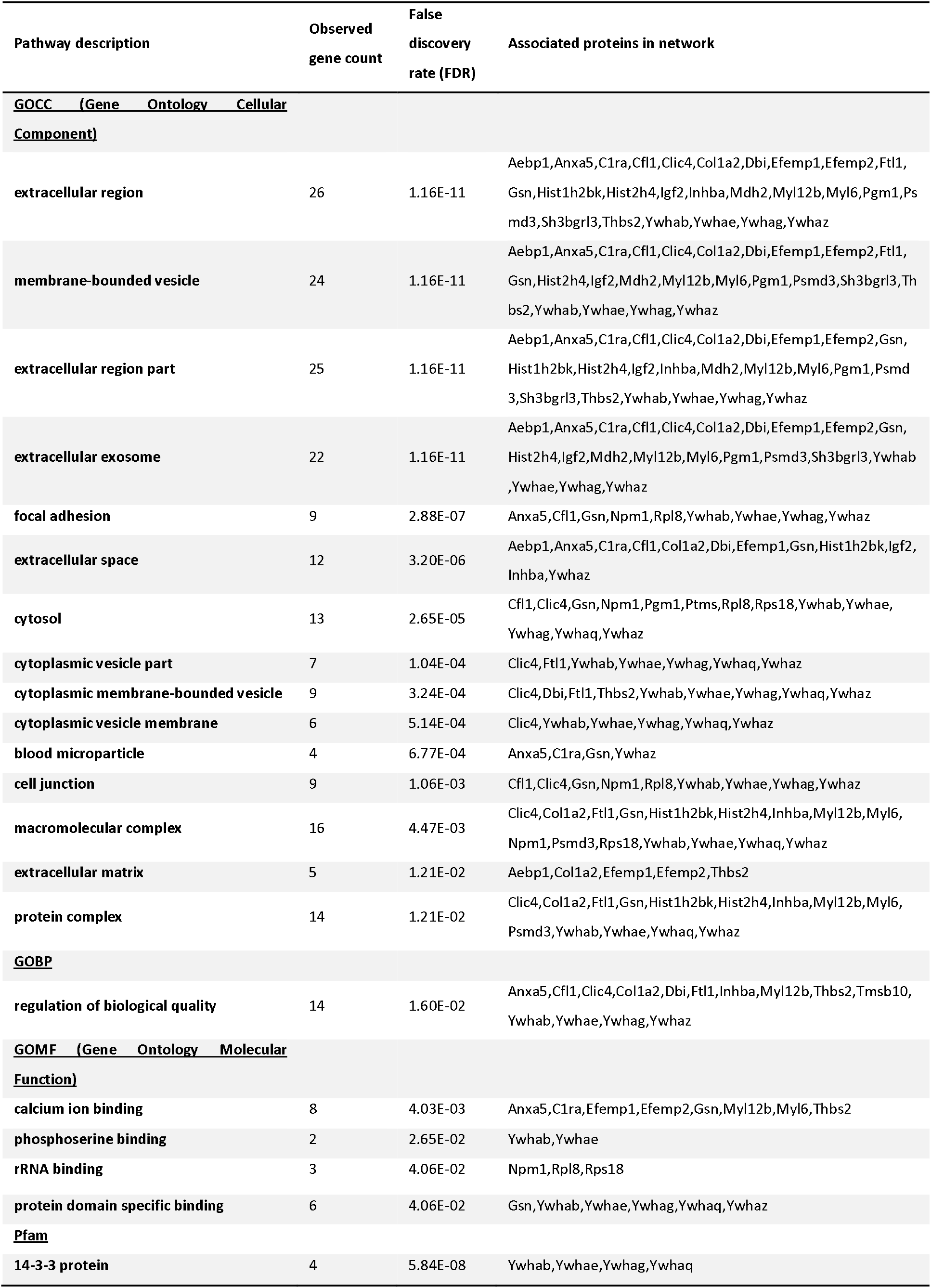
Functional enrichments in network using String DB with observed gene count out of a total of 34 genes and FDR cut-off of 2%.

**Figure S 1.**
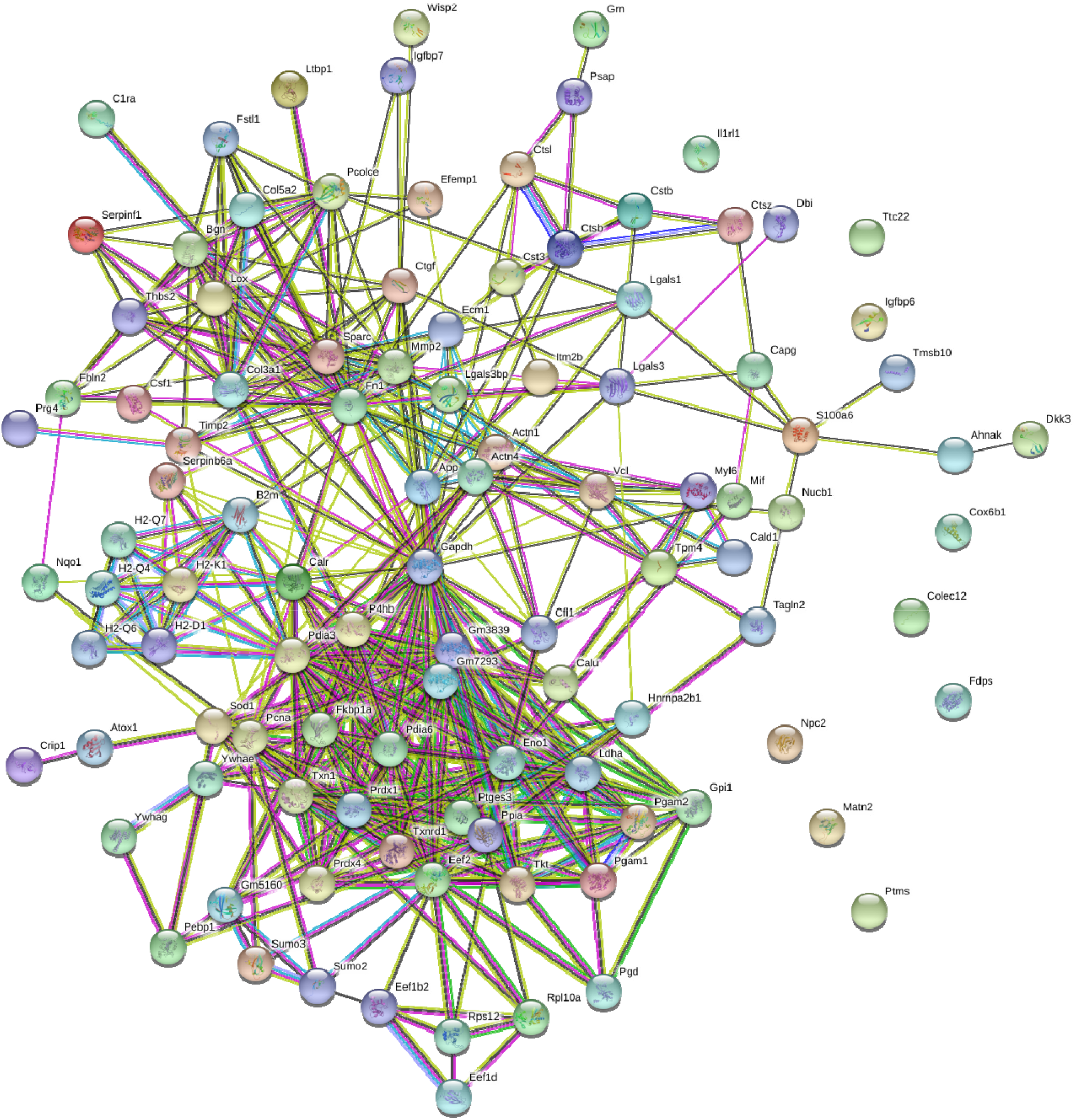
String DB network illustrating interactions between proteins in the osteocyte secretome, with significant degree of protein-protein interaction (p < 10^-16^)

**Figure S 2.**
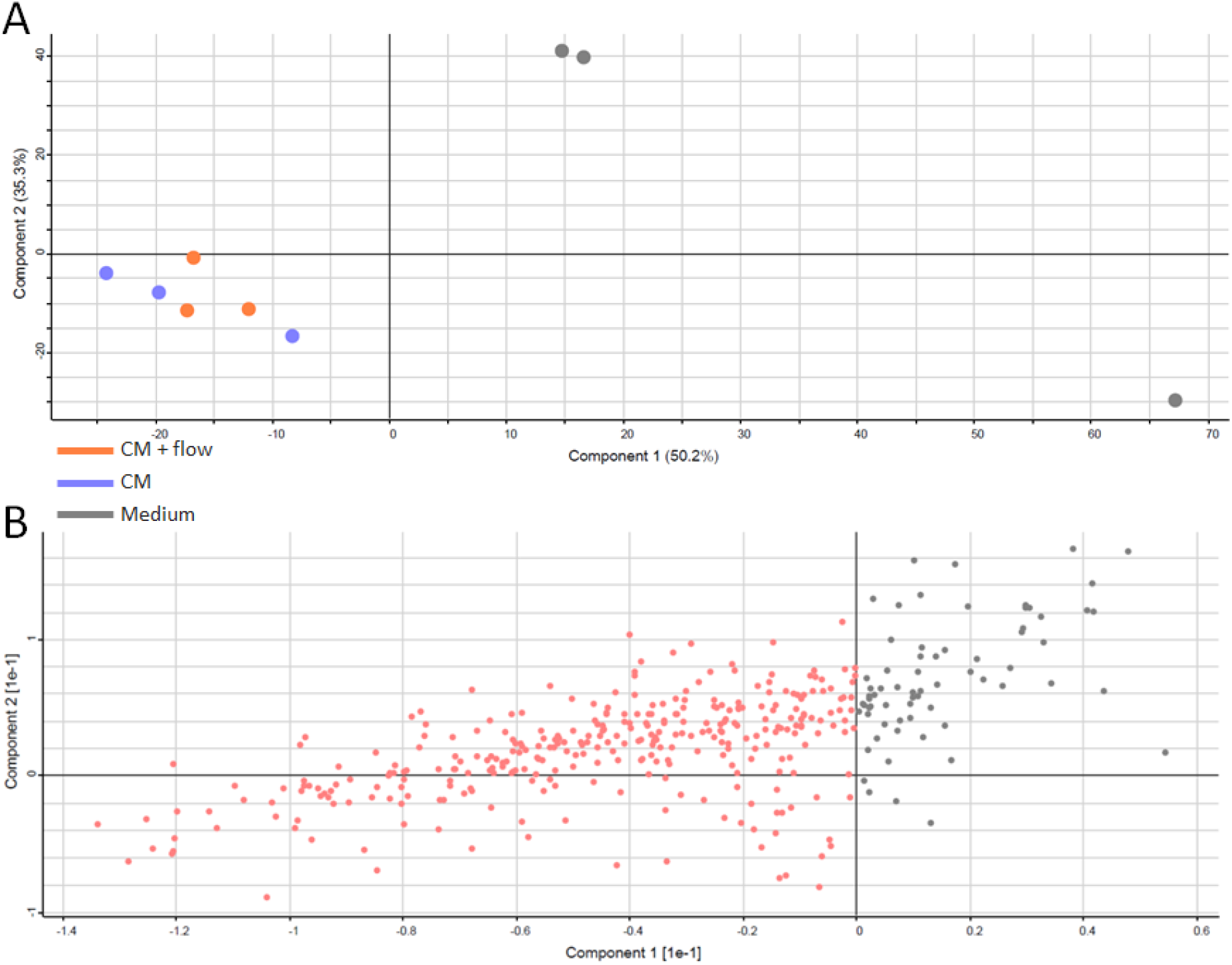
Principal Component Analysis (PCA) revealing the variance between the three experimental groups and indicating three main clusters of data (C). The proteins primarily driving the separation between the medium groups and the control groups are highlighted in red (D).

**Figure S 3.**
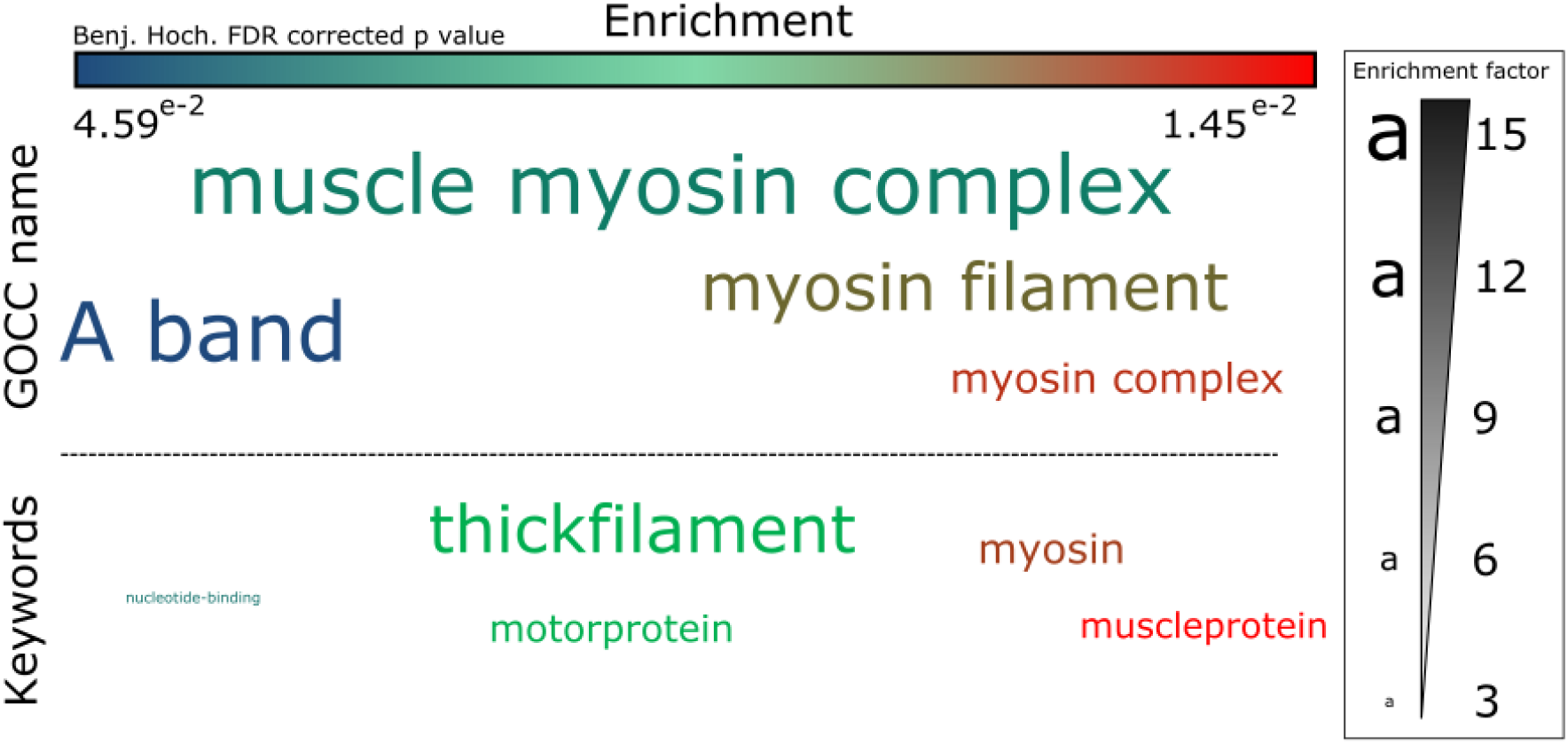
Enrichment analysis of GOCC terms and Uniprot keywords in proteins with greater expression in control medium samples, using Fisher’s exact test represented as a word cloud (D). The size of the word represents enrichment of terms, while colour represents FDR corrected p value. All terms with a minimum 0.05 FDR corrected p value were included.

**Table S 4.**
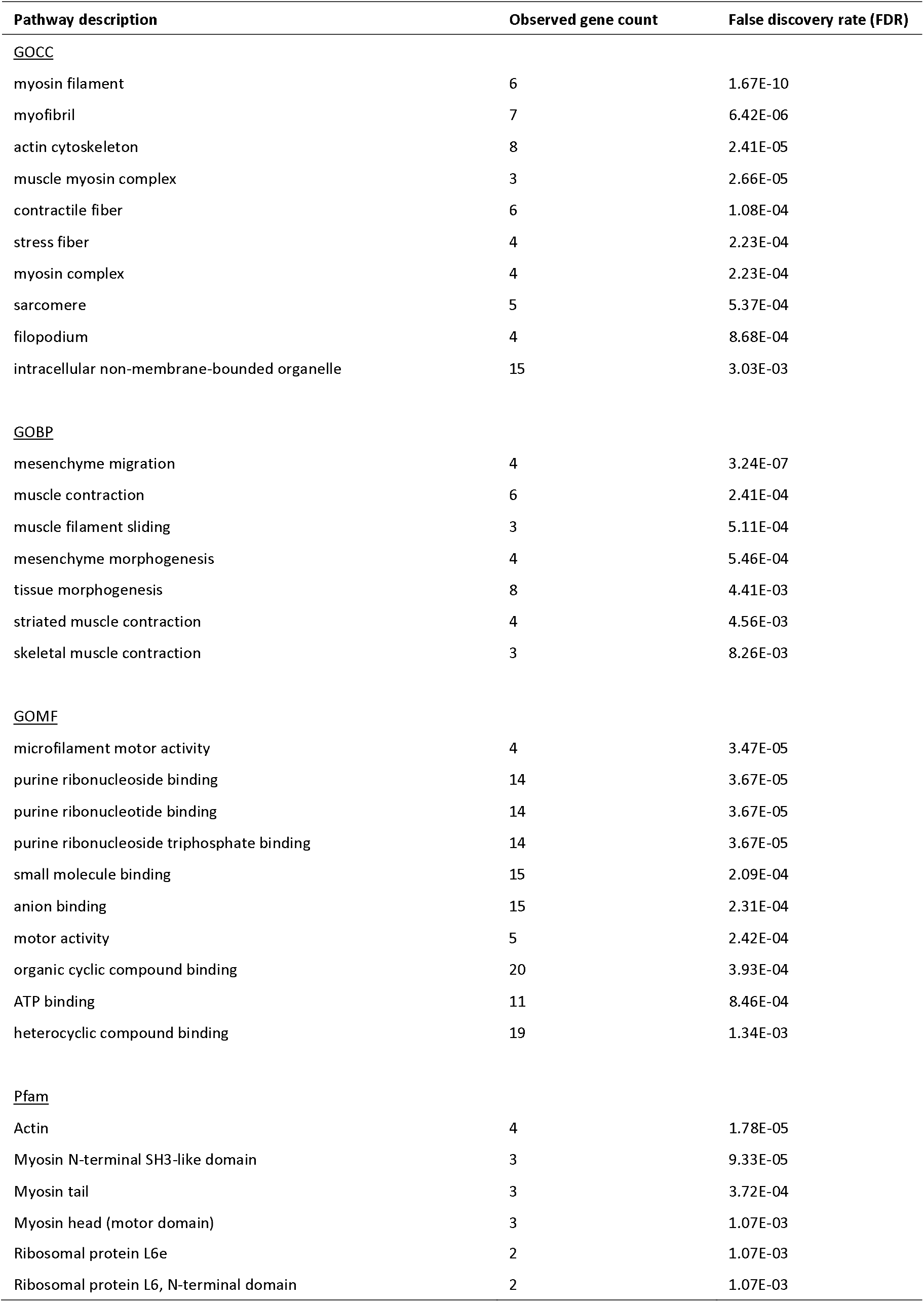
Functional enrichments in Medium proteins using String DB, with observed gene count out of a total 35 proteins with an FDR cut-off of 2%.

**Figure S 4.**
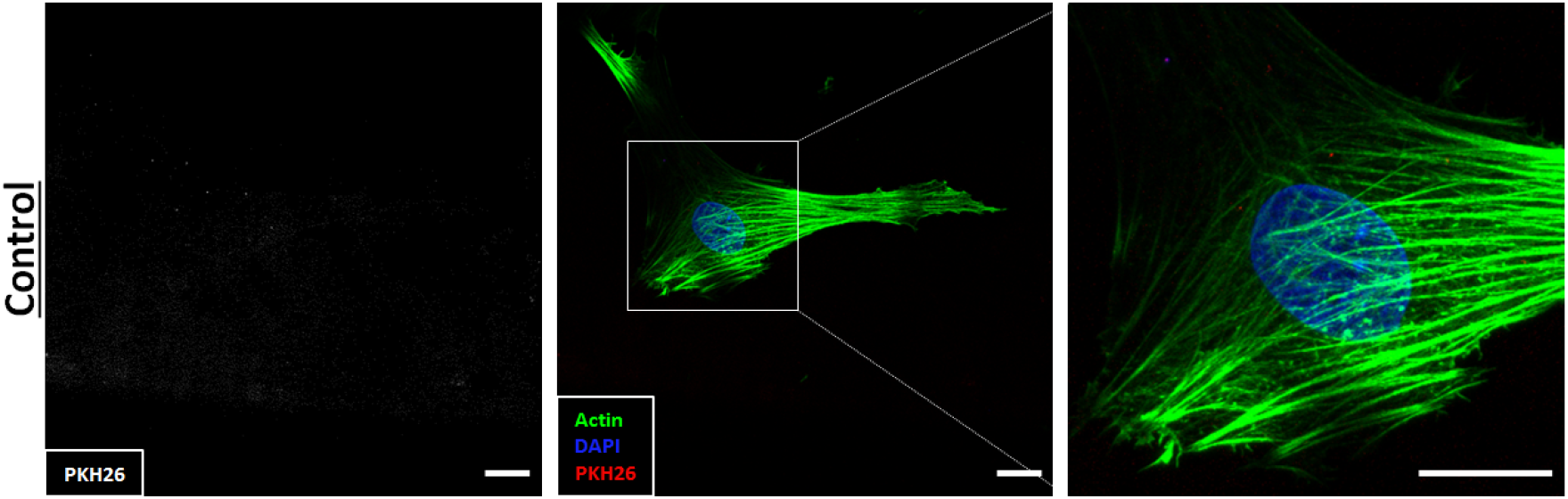
Control samples with no EVs and PKH26 staining demonstrating minimal unspecific fluorescence (Scale = 10μm).

## Acknowledgments

The authors would like to acknowledge funding from European Research Council (ERC) Starting and Proof of Concept Grant (336882 & 825905), Science Foundation Ireland (SFI) Support Grant SFI 13/ERC/L2864 and Irish Research Council Postgraduate Scholarship (GOIPG/2014/493)

## References

1. Dallas, S.L., M. Prideaux, and L.F. Bonewald, The osteocyte: An endocrine cell… and more. Endocrine Reviews, 2013. 34(5): p. 658–690.

2. Bonewald, L.F., Osteocyte biology: its implications for osteoporosis. J Musculoskelet Neuronal Interact, 2004. 4(1): p. 101–4.

3. Qiu, S., et al., Reduced iliac cancellous osteocyte density in patients with osteoporotic vertebral fracture. J Bone Miner Res, 2003. 18(9): p. 1657–63.

4. Rachner, T.D., S. Khosla, and L.C. Hofbauer, Osteoporosis: now and the future. The Lancet, 2011. 377(9773): p. 1276–1287.

5. Bonewald, L.F., The Role of the Osteocyte in Bone and Nonbone Disease. Endocrinology and Metabolism Clinics of North America, 2017. 46(1): p. 1–18.

6. Zhou, J.Z., et al., Osteocytic connexin hemichannels suppress breast cancer growth and bone metastasis. Oncogene, 2016. 35(43): p. 5597–5607.

7. Bonewald, L.F., The amazing osteocyte. Journal of Bone and Mineral Research, 2011. 26(2): p. 229–238.

8. Schaffler, M.B. and O.D. Kennedy, Osteocyte Signaling in Bone. Current Osteoporosis Reports, 2012. 10(2): p. 118–125.

9. Taylor, A.F., et al., Mechanically stimulated osteocytes regulate osteoblastic activity via gap junctions. Am J Physiol Cell Physiol, 2007. 292(1): p. C545–52.

10. Brady, R.T., F.J. O’Brien, and D.A. Hoey, Mechanically stimulated bone cells secrete paracrine factors that regulate osteoprogenitor recruitment, proliferation, and differentiation. Biochem Biophys Res Commun, 2015. 459(1): p. 118–23.

11. Hoey, D.A., D.J. Kelly, and C.R. Jacobs, A role for the primary cilium in paracrine signaling between mechanically stimulated osteocytes and mesenchymal stem cells. Biochemical and Biophysical Research Communications, 2011. 412(1): p. 182–187.

12. Tan, S.D., et al., Osteocytes subjected to fluid flow inhibit osteoclast formation and bone resorption. Bone, 2007. 41(5): p. 745–751.

13. You, L., et al., Osteocytes as Mechanosensors in the Inhibition of Bone Resorption Due to Mechanical Loading Bone, 2008. 42(1): p. 172–179.

14. Park, D, et al., Endogenous bone marrow MSCs are dynamic, fate-restricted participants in bone maintenance and regeneration. Cell Stem Cell, 2012. 10(3): p. 259–72.

15. McClung, M.R., Sclerostin antibodies in osteoporosis: latest evidence and therapeutic potential. Therapeutic advances in musculoskeletal disease, 2017. 9(10): p. 263–270.

16. Chen, W., et al., Gene expression patterns of osteocyte-like MLO-Y4 cells in response to cyclic compressive force stimulation. Cell Biology International, 2010. 34(5): p. 425–432.

17. Wasserman, E., et al., Differential load-regulated global gene expression in mouse trabecular osteocytes. Bone, 2013. 53(1): p. 14–23.

18. Govey, P.M., et al., Integrative transcriptomic and proteomic analysis of osteocytic cells exposed to fluid flow reveals novel mechano-sensitive signaling pathways. J Biomech, 2014. 47(8): p. 1838–45.

19. Yanez-Mo, M., et al., Biological properties of extracellular vesicles and their physiological functions. J Extracell Vesicles, 2015. 4: p. 27066.

20. Davies, O.G., et al., Annexin-enriched osteoblast-derived vesicles act as an extracellular site of mineral nucleation within developing stem cell cultures. Sci Rep, 2017. 7(1): p. 12639.

21. Morhayim, J., et al., Paracrine Signaling by Extracellular Vesicles via Osteoblasts. Current Molecular Biology Reports, 2016. 2(1): p. 48–55.

22. Cui, Y., et al., Exosomes derived from mineralizing osteoblasts promote ST2 cell osteogenic differentiation by alteration of microRNA expression. FEBS Lett, 2016. 590(1): p. 185–92.

23. Li, Q., et al., Extracellular vesicle-mediated bone metabolism in the bone microenvironment. J Bone Miner Metab, 2018. 36(1): p. 1–11.

24. Sato, M., et al., Circulating osteocyte-derived exosomes contain miRNAs which are enriched in exosomes from MLO-Y4 cells. Biomedical Reports, 2017. 6(2): p. 223–231.

25. Qin, Y., et al., Myostatin inhibits osteoblastic differentiation by suppressing osteocyte-derived exosomal microRNA-218: A novel mechanism in muscle-bone communication. J Biol Chem, 2017. 292(26): p. 11021–11033.

26. Gimona, M., et al., Manufacturing of Human Extracellular Vesicle-Based Therapeutics for Clinical Use. Int J Mol Sci, 2017. 18(6).

27. Cappariello, A., et al., Osteoblast-derived extracellular vesicles are biological tools for the delivery of active molecules to bone. J Bone Miner Res, 2017.

28. Whitham, M., et al., Extracellular Vesicles Provide a Means for Tissue Crosstalk during Exercise. Cell Metabolism, 2018. 27(1): p. 237–251.e4.

29. Bab, I., et al., Histone H4-related osteogenic growth peptide (OGP): A novel circulating stimulator of osteoblastic activity. EMBO Journal, 1992. 11(5): p. 1867–1873.

30. Southwick, F.S., Gelsolin and ADF/cofilin enhance the actin dynamics of motile cells. Proceedings of the National Academy of Sciences, 2000. 97(13): p. 6936.

31. Thouverey, C., et al., Proteomic characterization of biogenesis and functions of matrix vesicles released from mineralizing human osteoblast-like cells. Journal of Proteomics, 2011. 74(7): p. 1123–1134.

32. Altschul, S.F., et al., Gapped BLAST and PSI-BLAST: a new generation of protein database search programs. Nucleic Acids Res, 1997. 25(17): p. 3389–402.

33. Kent, W.J., et al., The human genome browser at UCSC. Genome Res, 2002. 12(6): p. 996–1006.

34. Turner, C.H., et al., Recruitment and proliferative responses of osteoblasts after mechanical loading in vivo determined using sustained-release bromodeoxyuridine. Bone, 1998. 22(5): p. 463–9.

35. Chen, J.C., et al., Mechanical signals promote osteogenic fate through a primary cilia-mediated mechanism. FASEB J, 2016. 30(4): p. 1504–11.

36. Vezeridis, P.S., et al., Osteocytes subjected to pulsating fluid flow regulate osteoblast proliferation and differentiation. Biochem Biophys Res Commun, 2006. 348(3): p. 1082 8.

37. Schaffler, M.B., et al., Osteocytes: master orchestrators of bone. Calcif Tissue Int, 2014. 94(1): p. 5–24.

38. Govey, P.M., A.E. Loiselle, and H.J. Donahue, Biophysical regulation of stem cell differentiation. Curr Osteoporos Rep, 2013. 11(2): p. 83–91.

39. Liu, Y., et al., Homodimerization of Ror2 tyrosine kinase receptor induces 14-3-3(beta) phosphorylation and promotes osteoblast differentiation and bone formation. Mol Endocrinol, 2007. 21(12): p. 3050–61.

40. Priam, S., et al., Identification of soluble 14-3-3 as a novel subchondral bone mediator involved in cartilage degradation in osteoarthritis. Arthritis Rheum, 2013. 65(7): p. 1831–42.

41. Rodan, G.A. and A.R. Rodan, The family of osteoblast transcription factors is growing. IBMS BoneKEy, 2005. 2(10): p. 12–15.

42. Shen, J., et al., Transcriptional induction of the osteocalcin gene during osteoblast differentiation involves acetylation of histones H3 and H4. Molecular Endocrinology, 2003. 17(4): p. 743–756.

43. Dudakovic, A., et al., Histone deacetylase inhibition promotes osteoblast maturation by altering the histone H4 epigenome and reduces Akt phosphorylation. J Biol Chem, 2013. 288(40): p. 28783–91.

44. Paino, F., et al., Histone Deacetylase Inhibition with Valproic Acid Downregulates Osteocalcin Gene Expression in Human Dental Pulp Stem Cells and Osteoblasts: Evidence for HDAC2 Involvement. Stem Cells (Dayton, Ohio), 2014. 32(1): p. 279–289.

45. Pigossi, S.C., et al., Role of Osteogenic Growth Peptide (OGP) and OGP(10–14) in Bone Regeneration: A Review. International Journal of Molecular Sciences, 2016. 17(11): p. 1885.

46. Haut Donahue, T.L., et al., Annexin V disruption impairs mechanically induced calcium signaling in osteoblastic cells. Bone, 2004. 35(3): p. 656–63.

47. Genetos, D.C., et al., Impaired osteoblast differentiation in annexin A2- and -A5-deficient cells. PLoS One, 2014. 9(9): p. e107482.

48. Bornstein, P., et al., Thrombospondin 2 modulates collagen fibrillogenesis and angiogenesis. J Investig Dermatol Symp Proc, 2000. 5(1): p. 61–6.

49. Hankenson, K.D., et al., Increased marrow-derived osteoprogenitor cells and endosteal bone formation in mice lacking thrombospondin 2. Journal of Bone and Mineral Research, 2000. 15(5): p. 851–862.

50. Miedel, E., et al., Disruption of thrombospondin-2 accelerates ischemic fracture healing. Journal of Orthopaedic Research, 2013. 31(6): p. 935–943.

51. Morrell, A.E., et al., Mechanically induced Ca2+ oscillations in osteocytes release extracellular vesicles and enhance bone formation. Bone Research, 2018. 6(1): p. 6.

52. Morhayim, J., et al., Proteomic signatures of extracellular vesicles secreted by nonmineralizing and mineralizing human osteoblasts and stimulation of tumor cell growth. FASEB J, 2015. 29(1): p. 274–85.

53. Huang, C.-C., et al., Exosomes as Biomimetic Tools for Stem Cell Differentiation: Applications in Dental Pulp Tissue Regeneration. Biomaterials, 2016. 111: p. 103–115.

54. Xie, H., et al., Extracellular Vesicle-functionalized Decalcified Bone Matrix Scaffolds with Enhanced Pro-angiogenic and Pro-bone Regeneration Activities. Sci Rep, 2017. 7: p. 45622.

55. Diomede, F., et al., Three-dimensional printed PLA scaffold and human gingival stem cell-derived extracellular vesicles: a new tool for bone defect repair. Stem Cell Research & Therapy, 2018. 9: p. 104.

56. Davies, O.G., et al., Annexin-enriched osteoblast-derived vesicles act as an extracellular site of mineral nucleation within developing stem cell cultures. Scientific Reports, 2017. 7(1).

57. Golub, E.E., Role of matrix vesicles in biomineralization. Biochim Biophys Acta, 2009. 1790(12): p. 1592–8.

58. Muhlrad, A., et al., Inorganic phosphate regulates the binding of cofilin to actin filaments. Febs j, 2006. 273(7): p. 1488–96.

59. Tavafoghi, M. and M. Cerruti, The role of amino acids in hydroxyapatite mineralization. Journal of The Royal Society Interface, 2016. 13(123).

60. Cheng, B., et al., PGE(2) is essential for gap junction-mediated intercellular communication between osteocyte-like MLO-Y4 cells in response to mechanical strain. Endocrinology, 2001. 142(8): p. 3464–73.

61. Zhao, S., et al., MLO-Y4 osteocyte-like cells support osteoclast formation and activation. Journal of Bone and Mineral Research, 2002. 17(11): p. 2068–2079.

62. Shah, K.M., et al., Osteocyte isolation and culture methods. BoneKEy reports, 2016. 5: p. 838–838.

63. Kato, Y., et al., Establishment of an Osteocyte-like Cell Line, MLO-Y4. Journal of Bone and Mineral Research, 1997. 12(12): p. 2014–2023.

64. Rosser, J. and L.F. Bonewald, Studying osteocyte function using the cell lines MLO-Y4 and MLO-A5. Methods Mol Biol, 2012. 816: p. 67–81.

65. Stavenschi, E., M.-N. Labour, and D.A. Hoey, Oscillatory fluid How induces the osteogenic lineage commitment of mesenchymal stem cells: The effect of shear stress magnitude, frequency, and duration. Journal of Biomechanics, 2017. 55: p. 99–106.

66. Cox, J. and M. Mann, MaxQuant enables high peptide identification rates, individualized p.p.b.-range mass accuracies and proteome-wide protein quantification. Nature Biotechnology, 2008. 26(12): p. 1367–1372.

67. Tyanova, S., T. Temu, and J. Cox, The MaxQuant computational platform for mass spectrometry-based shotgun proteomics. Nature Protocols, 2016. 11(12): p. 2301–2319.

68. Cox, J., et al., Andromeda: a peptide search engine integrated into the MaxQuant environment. J Proteome Res, 2011. 10(4): p. 1794–805.

69. Cox, J., et al., Accurate Proteome-wide Label-free Quantification by Delayed Normalization and Maximal Peptide Ratio Extraction, Termed MaxLFQ. Molecular & Cellular Proteomics: MCP, 2014. 13(9): p. 2513–2526.

70. Tyanova, S., et al., The Perseus computational platform for comprehensive analysis of (prote)omics data. 2016. 13(9): p. 731–40.

71. Szklarczyk, D., et al., STRING v10: Protein-protein interaction networks, integrated over the tree of life. Nucleic Acids Research, 2015. 43(D1): p. D447–D452.

